# First-principles model of optimal translation factors stoichiometry

**DOI:** 10.1101/2021.04.02.438287

**Authors:** Jean-Benoît Lalanne, Gene-Wei Li

## Abstract

Enzymatic pathways have evolved uniquely preferred protein expression stoichiometry in living cells, but our ability to predict the optimal abundances from basic properties remains underdeveloped. Here we report a biophysical, first-principles model of growth optimization for core mRNA translation, a multi-enzyme system that involves proteins with a broadly conserved stoichiometry spanning two orders of magnitude. We show that a parsimonious flux model constrained by proteome allocation is sufficient to predict the conserved ratios of translation factors through maximization of ribosome usage The analytical solutions, without free parameters, provide an interpretable framework for the observed hierarchy of expression levels based on simple biophysical properties, such as diffusion constants and protein sizes. Our results provide an intuitive and quantitative understanding for the construction of a central process of life, as well as a path toward rational design of pathway-specific enzyme expression stoichiometry.

## Introduction

A universal challenge faced by both evolution and synthetic pathway creation is to optimize the cellular abundance of proteins. This abundance optimization problem is not only multidimensional – often involving several proteins participating in the same pathway – but also under systems-wide constraints, such as limited physical space [30] and finite nutrient inputs [70]. The complexity of this problem has prevented rational design of protein expression for pathway engineering [23]. Fundamentally, being able to predict the optimal and observed cellular protein abundances from their individual properties would reflect an ultimate understanding of molecular and systems biology.

Evolutionary comparison of gene expression across microorganisms suggests that basic principles governing the optimization problem may exist. We recently reported broad conservation of relative protein synthesis rates within individual pathways, even under circumstances in which the relative transcription and translation rates for the homologous enzymes have dramatically diverged across species [34]. Moreover, distinct proteins that evolved convergently towards the same biological function also displayed the same stoichiometry of protein synthesis in their respective species. These results suggest that the determinants of optimal in-pathway protein stoichiometry are likely modular and independent of detailed biochemical or physiological properties that differ across clades. However, the precise nature of such determinants remains unknown.

Translation of mRNA into proteins is a central pathway required for cell growth and therefore serves as an entry point for establishing a quantitative model of optimal in-pathway stoichiometry. In addition to ribosomes which have well-coordinated synthesis of subunits [50], nearly 100 protein factors are involved in facilitating ribosome assembly and translation initiation, elongation, and termination [14, 44, 57]. The intracellular abundances of these factors vary over 100-fold [38, 54], and their ratios are often maintained in different growth conditions and across different species [34]. As a group, the total amount of translation-related proteins per cell mass scales linearly with growth rate in most conditions [13, 60, 62]. This relationship is considered a bacterial ‘growth law’ and can be understood as a consequence of producing just enough ribosomes necessary to duplicate the proteome at a given growth rate. However, the relative stoichiometry among translation factors is less understood. Notable exceptions are the elongation factor Tu (EF-Tu) [17, 30] and more recently elongation factor Ts (EF-Ts) [21], which have been suggested to maximize translational flux per unit proteome. The selective pressure on the expression of other factors, which can be much more lowly expressed and often assumed to be non-limiting, remains to be determined.

Here we sought to derive an intuitive model to understand the quantitative abundance hierarchy (Fig. 1B) among the core translation factors (TFs), which have well-characterized functions (Table 1, schematic in Fig. 1A). Our goal is not to exhaustively model the heterogeneous movement of ribosomes on the transcriptome [16, 56, 64, 65] or to include as many details of the underlying molecular steps as possible [21, 66]. Instead, we coarse-grained global translation into a cycle that consists of sequential steps with interconnected fluxes that depend on core TFs concentrations. At steady-state cell growth, all individual fluxes are matched and the overall rate of ribosomes completing the full translation cycle is proportional to cell growth. By solving for the maximum flux under previously defined proteome allocation constraints [17, 21, 62], we obtained analytical solutions for the optimal factor concentrations, which agree well with the observed values. The ratios of optimal concentrations depend only on simple biophysical parameters that are broadly conserved across species. For instance, elongation factor EF-G is predicted to be more abundant than initiation and termination TFs by a multiplicative factor of 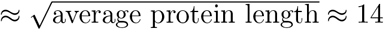, whereas EF-Tu is predicted to be more abundant than EF-G by a factor of 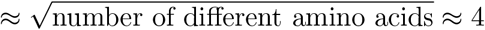. These results, arising from the optimization procedure and generic properties of the translation cycle, provide rationales for the order-of-magnitude expression of these important enzymes.

**Table 1:** Brief description of the function of core translation factors considered. For reviews of mRNA translation, see [12, 57].

| Step | Factor | Function |
| --- | --- | --- |
| Initiation | IF1 | Initiation factor 1: binds to 30S ribosome subunits to facilitate initiator tRNA binding [20, 36]. |
| Initiation | IF2 | Initiation factor 2: ribosome-dependent GTPase interacting with 30 ribosome subunits, ensures correct binding of initiator tRNAs [20, 36]. |
| Initiation | IF3 | Initiation factor 3: prevents premature docking of 50S ribosomal subunits [20, 36]. |
| Elongation | EF-Tu | Elongation factor Tu: binds to charged tRNAs to form ternary complexes, brings charged tRNAs to empty ribosome A sites. [1, 2, 68] |
| Elongation | aaRS | tRNA synthetases: charge tRNAs with cognate amino acids [22, 51]. |
| Elongation | EF-G | Elongation factor G: catalyzes translocation steps of the ribosome after peptide bond formation [1, 2]. |
| Elongation | EF-Ts | Elongation factor Ts: nucleotide exchange factor for EF-Tu [1, 2]. |
| Termination | RF1/RF2 | Peptide chain release factors 1 and 2: recognize stop codon and hydrolyze the completed protein. RF1 recognizes UAA, UAG, and RF2 UAA, UGA [8]. |
| Termination | RF4 | Ribosome recycling factor: catalyzes the dissociation of ribosome subunits following peptide chain release in translation termination [8]. |

**Figure 1:**
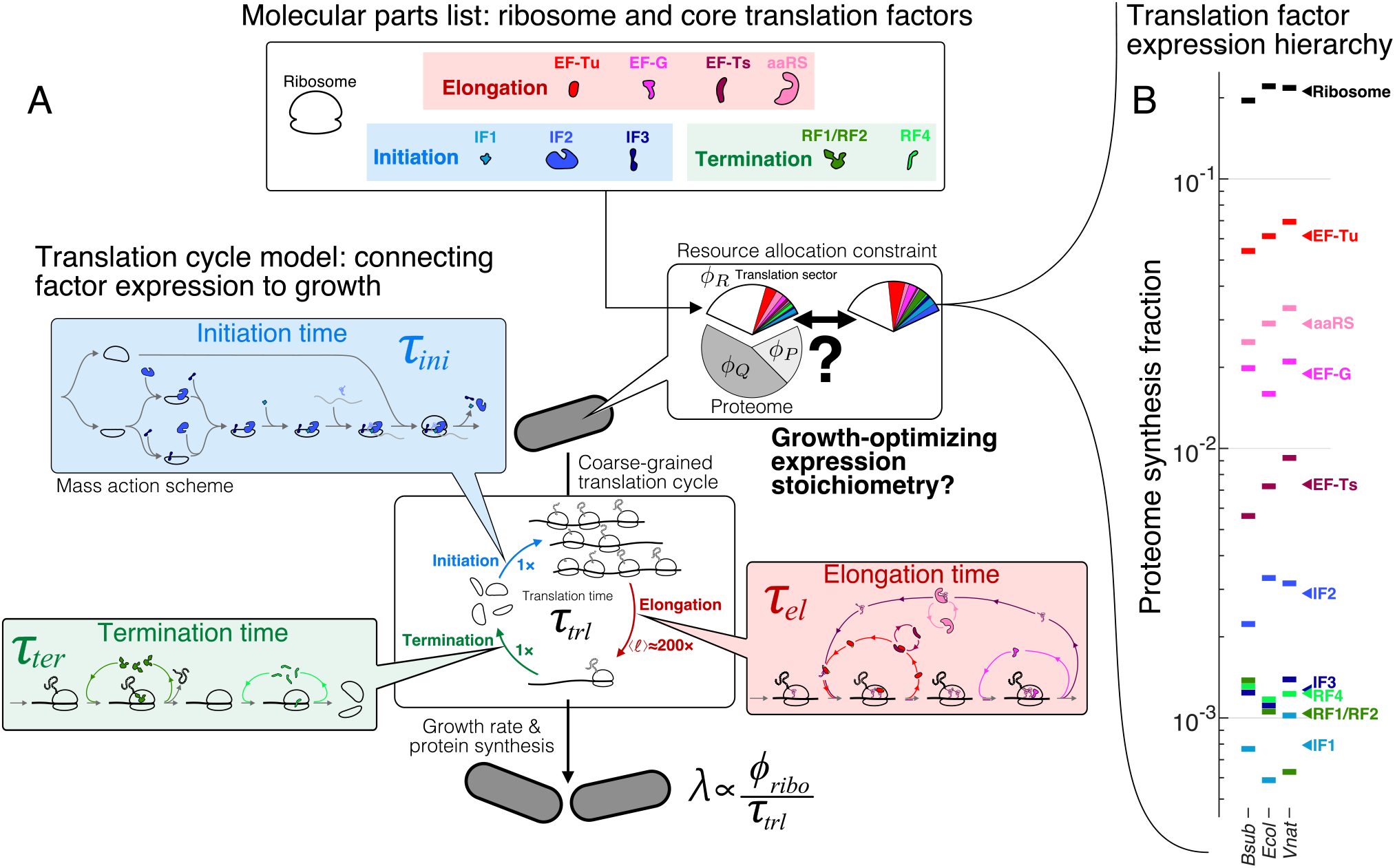
The hierarchy of mRNA translation factor expression stoichiometry. (A) Multiscale model relating translation factor expression to growth rate. The growth rate *λ* is directly proportional to the ribosome content (*ϕ* _*ribo*_) in the cell and inversely proportional to the average time to complete the translation cycle τ _*trl*_, consisting of the sum of the initiation (τ_*ini*_), elongation (τ_*el*_), and termination (τ_*ter*_) times. Each of these reaction times are determined by the translation factor abundances. On average, the elongation step is repeated around ⟨*ℓ* ⟩ ≈ 200 × to complete a full protein, compared to 1× for initiation and termination. Our framework of flux optimization under proteome allocation constraint addresses what ribosome and translation factor abundances maximize growth rate. (B) Measured expression hierarchy of bacterial mRNA translation factors, conserved across evolution. Horizontal bars mark the proteome synthesis fractions as measured by ribosome profiling [34] (equal to the proteome fraction by weight for a stable proteome) for key mRNA translation factors in *B. subtilis* (*Bsub*), *E. coli* (*Ecol*), and *V. natriegens* (*Vnat*) and are color-coded according to the protein (or group of proteins) specified. Triangles (◂) on the right indicate the mean synthesis fraction of the protein in the three species. See Table 1 for a short description of the translation factors considered.

## Results

### Problem statement and model formulation

Our overall goal is to determine the growth-optimizing proteome allocation for the ribosome and various translation factors. Conceptually, varying TF concentrations has two opposing effects on cell proliferation. At the biochemical level, high TF expression can facilitate growth by allowing more efficient usage of ribosomes. At the systems level, taking into account the constraint of a finite proteomic space, high TF expression can nonetheless limit growth by reducing the number of ribosomes produced. The tradeoffs between various TFs and ribosomes create a multidimensional optimization problem.

We solve this multidimensional problem by treating translation as a dynamical system, in which ribosomes cycle through initiation, elongation, and termination. The resulting flux drives cell growth. During steady-state growth, every interlocked step of the translation cycle must have the same ribosome flux that is specified by the growth rate. We show that at the growth optimum, concentrations for distinct TFs can be solved independently. The resulting analytical solutions can be expressed in terms of the growth rate and simple biophysical parameters.

### Cell growth driven by TF-dependent ribosome flux

To describe the biochemical effects of TF concentrations on cell growth, we first introduce a coarse-grained translation cycle time τ_*trl*_, or the time it takes for a ribosome to complete a typical cycle of protein synthesis (Fig. 1A), which consists of three sequential steps: initiation (“*ini*”), elongation (“*el*”), and termination (“*ter*”). Each of these steps is catalyzed by multiple TFs. The full translation cycle time is then sum of ribosome transit times at the three steps (τ_*trl*_ = τ _*ini*_ + τ_*el*_+ τ_*ter*_), whose dependence on individual TF concentrations can be quantitatively described through mass action kinetic schemes (schematically depicted in Fig. 1A, see Supplement and examples below). We express TF concentrations in units of proteome fractions (dry mass fraction of a specified protein to the full proteome), denoted by *ϕ* [62] (Supplement, section S1.2). Using this notation, the translation cycle time τ_*trl*_ is a decreasing function of various TFs concentrations ({*ϕ*_*TF,i*_}).

In addition to its dependency on TF concentrations, the translation cycle time provides a bridge between the cell growth rate and ribosome concentration. In steady-state growth [13, 47, 62], the growth rate of cells and of their protein content (total number of proteins) must be identical, denoted here as *λ*, as a result of the constant average cellular composition. The protein content grows at a rate determined by the flux of ribosomes completing the translation cycle (*N*_*ribo*_*/*τ_*trl*_, where *N*_*ribo*_ is the number of ribosomes per cell) divided by the total number of proteins *N*_*P*_ per cell, i.e., *λ* = *N*_*ribo*_*/*τ_*trl*_*N*_*P*_. Rescaling the ribosome number to the mass fraction of proteome for ribosomes (*ϕ*_*ribo*_) yields

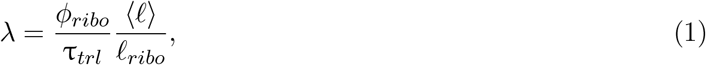

where *ℓ*_*ribo*_ is the number of amino acids in ribosomal proteins and ⟨*ℓ*⟩ is the average length among all proteins, weighted by expression levels (Methods, section S1). The rescaling factor (*ℓ*_*ribo*_*/*⟨*ℓ*⟩≈ 7300 *a*.*a*.*/*200 *a*.*a*. = 36.5) is approximately constant across growth conditions (section S1). This equation establishes how TF concentrations affect the growth rate biochemically via τ_*trl*_.

We note that equation 1 is a generalized form of the bacterial growth law that relates the mass fraction of elongating ribosomes to growth rate (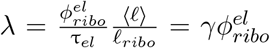, where *γ* is a rescaled translation elongation rate and 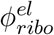 is the proteome fraction of actively translating ribosomes [13, 62, 63]). This classic growth law was derived by considering the steady-state flux of peptide bond formation by elongating ribosomes, whereas our model focuses on the flux of ribosomes that traverse the entire translation cycle, thereby allowing us to consider the effects of translation factors and ribosomes engaged in additional steps (initiation, elongation, and termination). For each step, equation 1 can be extended to show that the growth rate is similarly proportional to the mass fraction of the corresponding ribosomes divided by the transit time at that step (Supplement, section S2).

Steady-state growth thus imposes the requirement that the growth rate be inversely proportional to the translation cycle time and proportional to the number of ribosomes engaged in the translation cycle (equation 1). Inactive ribosomes, comprised of assembly intermediates, hibernating ribosomes, or otherwise non-functional ribosomes, have been found to constitute a small fraction of the total ribosome pool for fast growth [13, 39] and are not considered here. Based on equation 1, both increasing ribosome concentration and increasing TF concentrations (which decreases τ_*trl*_) can accelerate growth. However, production of ribosomes and TFs is subject to competition under a limited proteomic space, which we consider next.

### Optimization under proteome allocation constraint

To model the production cost tradeoff between TFs and ribosomes, we integrate the flux-based formulation above to the proteome allocation model that describes how global gene expression scales with the growth rate [17, 62]. In this coarse-grained model, the proteome is divided into an incompressible “sector” *ϕ*_*Q*_ that is independent of the growth rate, a metabolic protein sector *ϕ*_*P*_ that scales linearly with growth rate (*ϕ*_*P*_ := *λ/v*, where *v* is a phenomenological parameter that reflects nutrient quality), and a translation sector *ϕ*_*R*_. In our description, *ϕ*_*R*_ is taken as the summed proteome fractions for the ribosome and translation factors, i.e., *ϕ*_*R*_ := *ϕ*_*ribo*_+ Σ _*i*_ *ϕ*_*TF,i*_. Together, these sectors comprise the full proteome: 1 = *ϕ*_*Q*_ + *ϕ*_*P*_ + *ϕ*_*R*_. Under this framework, the proteomic tradeoff between ribosomes and TFs is then

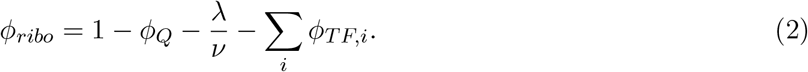

Equations 1 and 2, together with to the kinetic schemes for each step of the translation cycle, constitute the core of our model. Combining the biochemical effects (equation 1) and the systems-level constraints (equation 2) on TFs, we arrive at a self-contained relationship between growth and TF concentrations:

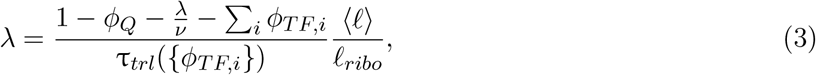

where we explicitly express τ_*trl*_ as a function of *ϕ*_*TF,i*_ to reflect the dependence of ribosome transit times on translation factor abundances. The above relationship (equation 3) allows us to ask: what is the stoichiometry of TFs, or partitioning of the translation sector, that maximizes the growth rate (Fig. 1A)?

The condition for the optimal TF abundances, i.e., the set of *ϕ*_*TF,i*_ that satisfies (*∂λ/∂ϕ*_*TF,i*_)^*^ = 0, can be readily obtained by considering the *ϕ*_*TF,i*_ as independent variables and taking the derivative of equation 3 with respect to a specified TF abundance, which yields

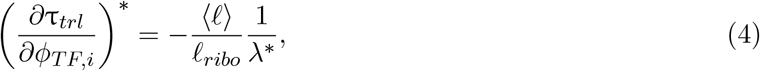

where the asterisk refers to the growth optimum, i.e., (*∂λ/∂ϕ*_*TF,i*_)^*^ = 0. Hence, under this framework, the TF abundances are growth-optimized when the sensitivity of the translation cycle time to changing the considered TF abundance (*∂*τ_*trl*_ */∂ϕ*_*TF,i*_) reaches a value determined solely by the growth rate and protein size factors.

Although equation 3 and the resulting optimization conditions (equation 4, one for every TF) corresponds to a coupled nonlinear system of multiple *ϕ*_*TF,i*_, substantial decoupling occurs at the optimal growth rate. In this situation, most *ϕ*_*TF,i*_ are only connected through the resulting growth rate. The optimization problem is then further simplified by the fact that the translation cycle consists of sequential and largely independent steps. The translation cycle time τ_*trl*_ corresponds to the sum of the coarse-grained initiation, elongation, and termination times, i.e., τ_*trl*_ = τ_*ini*_ + τ_*el*_ + τ_*ter*_. Given that eachTF is involved in a specific molecular step, the sensitivity matrix of these times to TF concentration is sparse: (*∂*τ_*j*_ */∂ϕ*_*TF,i*_)^*^ = 0 for most combinations of τ_*j*_ and *ϕ*_*TF,i*_. This lack of ‘cross-reactivity’ expresses that, for example, the initiation time τ_*ini*_ is unaffected by the tRNA synthetase concentration. This sparsity only occurs at the optimal expression levels, as the transit times typically depend on the growth rate (see an example in section S5.2) and *∂λ/∂ϕ*_*TF,i*_ ≠ 0 away from the optimum. The optimum condition for factor *i* then simplifies to:

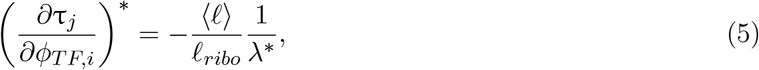

where *j* denotes the translation step(s) that TF_*i*_ participates in. This leads to simplifications that allow the system to be solved analytically in most cases: instead of solving the full system at once, individual reactions within the translation cycle can be considered in isolation. The resulting optimal concentrations are connected via the growth rate *λ*^*^. Interestingly, the optimal stoichiometry among most TFs is independent of *λ*^*^ if the reactions are diffusion-limited, as we show below.

### Case study: Translation termination

We first illustrate the process of solving for the optimal TF concentration for the relatively simple case of translation termination. The principles used here and the form of solutions provide conceptual guideposts for solving other steps of the translation cycle.

In bacteria, translation termination [8] consists of two distinct, sequential steps: (1) stop codon recognition and peptidyl-tRNA hydrolysis catalyzed by class I peptide chain release factors RF1 and RF2, followed by (2) dissociation of ribosomal subunits from the mRNA, i.e., ribosome recycling, catalyzed by RF4. We do not explicitly consider the additional factors (e.g., RF3 and EF-G) due to their lack of conservation or because they are non-limiting for this specific step (Supplement, section S5.1). RF1 and RF2 have the same molecular functions but recognize different stop codons [61]: RF1 recognizes stops UAA and UAG, whereas RF2 recognizes UAA and UGA. For simplicity, we describe here a scenario where RF1 and RF2 have no specificity towards the three stop codons, which allows us to combine them in a single factor (denoted RFI). The model is readily generalized, with similar results, to the case of the two RFs with their specificity towards the three stop codons (Supplement, section S5.3).

Under a coarse-grained description, the total ribosome transit time at termination τ_*ter*_ can be decomposed into a sum of peptide release time and ribosome recycling time. In the treatment below, we consider a regime of diffusion-limited reactions for simplicity. A full model with catalytic components can also be solved analytically (Supplement section S5.2, Fig. 2A). In the diffusion-limited regime (*k*_*cat*_ → ∞), the peptide release time and ribosome recycling time are inversely proportional to the corresponding TF concentrations:

**Figure 2:**
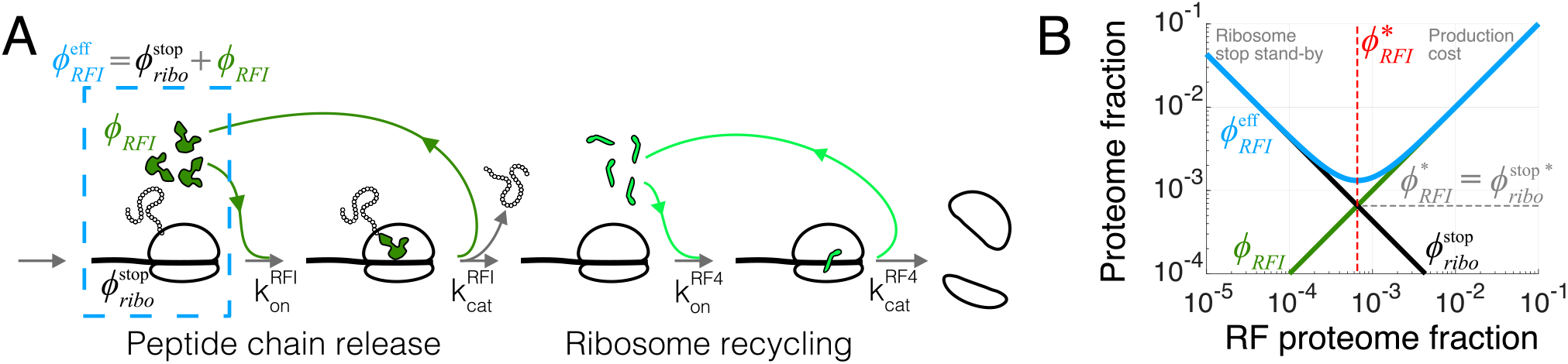
Case study with translation termination (A) Coarse-grained translation termination scheme. (B) Illustration of the minimization of effective proteome fraction corresponding to peptide chain release factors, leading to the equipartition principle.

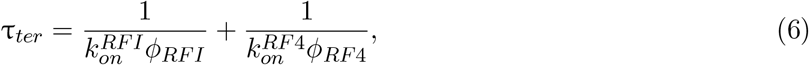

where the association rate constants 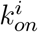 are rescaled by the factor’s sizes in proteome fraction units (Supplement, section S1.2). The above expression constitutes the solution of the mass action scheme for termination, connecting factor abundances to termination time.

The termination time (equation 6) can then be directly substituted into the optimality condition (equation 5) and solved in terms of *λ*^*^:

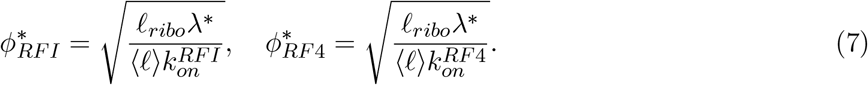

If the reactions are not diffusion-limited, an additional catalytic term ∝ *λ*^*^*/k*_*cat*_ is added to the minimally required levels above (Supplement, section S5.2). The square-root dependence in the optimal RF concentrations emerges from the 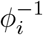 dependence of τ_*i*_, e.g., for ribosome recycling 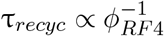, which becomes 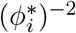 upon taking the derivative in the optimality condition (equation 5). The square root is then obtained by solving for 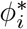. A similar square-root dependence has been noted in optimization of the ternary complex and tRNA abundances [6, 17]. As a result of the square-root, the optimal RF concentrations are weakly affected by biophysical properties such as the association rate constants and protein sizes. In the diffusion-limited regime above, the ratio of the optimal concentrations between RFI and RF4 is independent of the growth rate and only depends on the kinetics of binding.

As a side note, the expression for termination time τ_*ter*_ in equation 6 must be modified in a regime where ribosomes are frequently queued upstream of stop codons. This would occur if the termination rate were slow and approached initiation rates on mRNAs [7, 33]. In this regime, queues of ribosomes at stop codons would incur an additional time to terminate. In a general description, the resulting additional termination time can be absorbed in a queuing factor 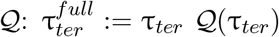 (Supplement, section S3 for derivation and discussion). The resulting nonlinearity would forbid the decoupling in the optimization procedure between RFI and RF4. Although absolute rates of termination are difficult to measure *in vivo*, translation on mRNAs is generally thought to be limited at the initiation step [36], and consistently, ribosome queuing at stop codons in bacteria is not usually observed (except under severe perturbations, e.g., [3, 27, 33, 42, 58]). In the physiological regime of fast termination, the queuing factor converges to 1, yielding simple solutions that depend only on biophysical parameters (equations 7).

### Equipartition between TF and corresponding ribosomes

The optimal TF concentrations (e.g., equation 7) can also be intuitively derived from another viewpoint. For each reaction in the translation cycle, we can define an effective proteome fraction allocated to that process, combining the proteome fractions of the corresponding TF and the ribosomes waiting at that specific step. As an example, for the case of peptide chain release factor (RFI) just treated, the effective proteome fraction includes the release factors and ribosomes with completed peptides waiting at stop codons (dashed box in Fig. 2A), i.e 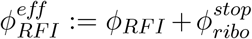. This effective proteome fraction corresponds to the total proteomic space associated to a TF in the context of the translation cycle.

During steady-state growth, the concentration of ribosomes waiting at any specific step of the translation cycle is equal to the total ribosome concentration multiplied by the ratio of the transit time of that step to the full cycle: e.g., here 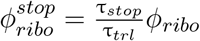, where 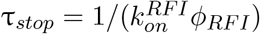 is the time to arrival of RFI. Using equation 1 for *ϕ*_*ribo*_, the effective proteome fraction satisfies:

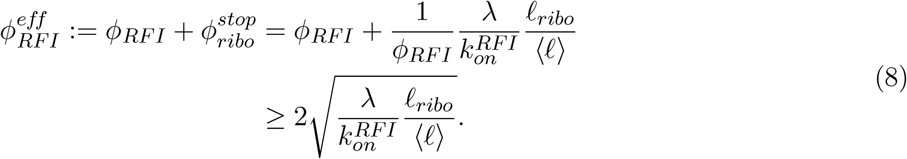

In the last line, we used the inequality of arithmetic and geometric means 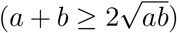 to obtain the minimum of the effective proteome fraction. The equality holds when the two proteome fractions are equal 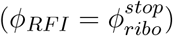, which provides the solution for optimal *ϕ*_*RFI*_ :

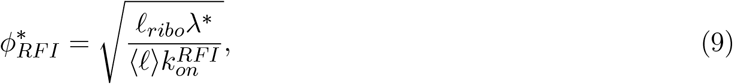

Hence, we recover equation 7 by minimizing the effective proteome fraction allocated to a given process in the translation cycle (the above argument applies to the free optimal concentration in the non-diffusion limited regime, see Supplement, section S5.2 for an example). From this perspective, optimization of the translation apparatus balances the production cost of the enzyme of interest with the improved efficiency of a having less ribosomes idle at that step, Fig. 2B. The optimal abundance corresponds to a point of equipartition: the proteome fraction of free cognate factors equals the proteome fraction of ribosomes waiting at the corresponding step (Fig. 2B).

### Case study: Ternary complex and tRNA cycle (EF-Tu and aaRS)

We next consider a more complex step of the translation cycle – elongation – and demonstrate that the optimality criteria (equation 5) can similarly provide simple analytical solutions in the physiologically relevant regime. Translation elongation involves multiple interlocked cycles and enzymes (EF-Tu, EF-G, EF-Ts, aminoacyl-tRNA synthetases (aaRS), and more). Our kinetic scheme for translation elongation, self-consistently coarse-grained to a single codon (derived in the Supplement, section S6.1), is shown in Fig. 3A. Charged tRNAs are brought to ribosomes through a ternary complex (TC), corresponding to a bound tRNA and EF-Tu. Following tRNA delivery and GTP hydrolysis, EF-Tu is released from the ribosome, and nucleotide exchange factor EF-Ts recycles EF-Tu back into the active pool, after which EF-Tu can bind a charged tRNA again and form another TC. At the ribosome, translocation to the next codon is catalyzed by EF-G, followed by release of uncharged tRNAs. Aminoacyl-tRNA synthetases then charge tRNAs to complete the elongation cycle.

**Figure 3:**
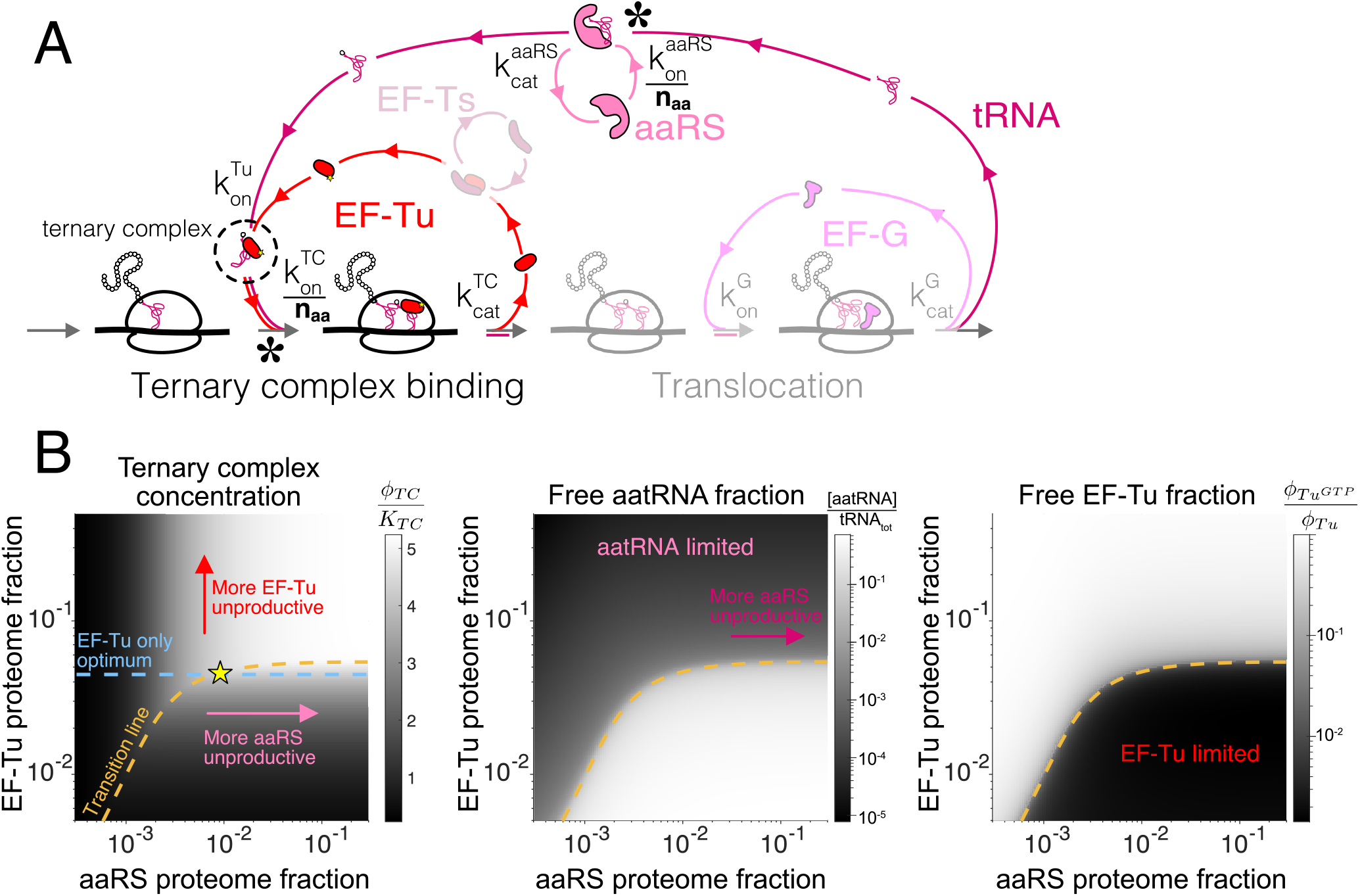
Case study with elongation factors (EF-Tu/aaRS) (A) Schematic of the translation elongation scheme, with the tRNA cycle, involving tRNA synthetases (aaRS) and EF-Tu. Reactions with * have the association rate constants rescaled by a factor of 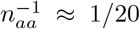 due to coarse-graining to a single codon model. Greyed out cycles (EF-Ts and EF-G) can be solved in isolation (Supplement, respectively sections S6.3.3 and S6.3.4). (B) Exploration of the aaRS/EF-Tu expression space from numerical solution of the elongation model (Supplement, section S6.3.5). The transition line (orange) marks the boundary between the EF-Tu limited and aaRS limited regimes. Left panel shows the ternary complex concentration (which is closely related to the elongation rate, equation 10). The ternary complex concentration is scaled by the dissociation constant *K*_*TC*_ to the ribosome A site (Supplement, section S6.3.5). Middle panel shows the free charged tRNA fraction. Right panel shows the free EF-Tu fraction (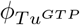 denotes the proteome fraction of EF-Tu GTP that can bind to charged tRNAs to form the ternary complex). The star marks the optimal solution, as described in the text. Middle panel shows the free charged tRNA fraction, and right panel the free EF-Tu fraction.

Similar to translation termination, the factor-dependent ribosome transit time through a single codon (τ_*aa*_) is comprised of two steps, corresponding to binding of the TC and EF-G, respectively (formal derivation and non-diffusion-limited regime in section S6.3.1):

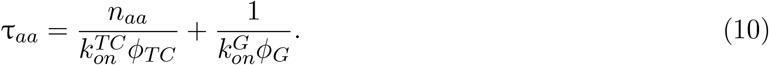

The coarse-grained total elongation time in our model is given by the factor-dependent transit time of a single codon above multiplied by the average protein sizes, i.e., τ_*el*_ = ⟨*ℓ*⟩(τ_*aa*_+τ_*ind*_). A factor-independent time τ_*ind*_ (e.g., peptidyl transfer), which does not come into play in our optimization framework, was added to account for additional steps making up the full elongation cycle. The rescaling of the TC association rate by *n*_*aa*_ arises as a result of our coarse-graining to a one-codon model, and reflects the codon specific interaction between the TC and the ribosome (Supplement, section S6.1). Note that the ternary complex concentration, *ϕ*_*TC*_, is a nonlinear function of the concentrations of all elongation factors (including *ϕ*_*G*_).

Despite the complexity of τ_*aa*_ as a function of the *ϕ*_*TF,i*_, the fact that all fluxes are equal in steady-state allows several steps to be isolated and solved separately (EF-Ts and EF-G, greyed out in Fig. 3A, respectively solved in Supplement, sections S6.3.3 and S6.3.4). For example, the approximate solution for optimal EF-G concentration parallels that for termination factors:

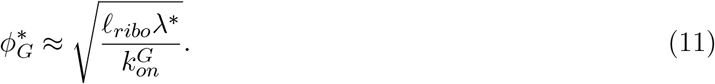

Importantly, the optimum for EF-G is larger than the optimum for RFs by a factor 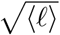, reflecting that a typical translation cycle requires ⟨*ℓ*⟩ steps catalyzed by EF-G and only one step for RFs (i.e., τ_*el*_ = ⟨*ℓ*⟩(τ_*aa*_ + τ_*ind*_) enters the optimality condition, equation 5, in contrast to τ_*ter*_ which is not multiplied by a scaling factor). The square root dependence arises here for the same reason as in the case of translation termination (derivative of *ϕ*^−1^).

In contrast to EF-G and EF-Ts, EF-Tu and aaRS cannot *a priori* be treated in isolation because the TC is composed of charged tRNAs and EF-Tu. A key property of the model simplifies the problem considerably. Following coarse graining, there is a separation of timescales between two classes of reactions in the elongation cycle: those that require codon specificity and those that do not. The first class includes aaRS binding to cognate tRNAs and TC binding to cognate codons (asterisks in Fig. 3A). The association rate constants of these codon-specific reactions are rescaled by a factor of 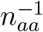, with *n* _*aa*_ ≈ 20 close the number of different amino acids (derivation and estimation of the coarse-graining parameter from experimental data in section S6.1). Heuristically, the rescaling arises from having *n*_*aa*_ parallel pathways whose rates are dictated by bimolecular reactions with respective components present at a concentration 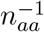 lower, with an overall rate reduction proportional to 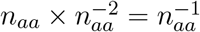. By comparison, the reactions that are codon-agnostic, such as binding of EF-Tu to charged tRNAs, are much faster.

The dependence of TC concentration (*ϕ*_*TC*_ in equation 10) on EF-Tu and aaRS concentrations can be solved based on this separation of timescales (Supplement, section S6.3.5) and the condition that EF-G and EF-Ts are at their respective optima. Here we consider an effective one-codon model and the corresponding sum of all aaRS abundances as *ϕ*_*aaRS*_. The EF-Ts portion of the EF-Tu cycle leads to an additive correction to the optimal EF-Tu solution and is omitted from the current discussion for simplicity, see section S6.3.5 for a full treatment.

As a result of the separation of timescales, the behavior of TC concentration in the two-dimensional EF-Tu/aaRS expression space is separated into two regimes: one where EF-Tu is limiting (free EF-Tu concentration near 0) for ternary complex formation, and the other where charged tRNAs are limiting (free charged tRNA concentration near 0). Specifically, in either limited regimes, increasing the abundance of the limiting component leads to stoichiometric increase in TC and therefore dictates TC concentration. The two regimes are separated by a transition region, whose width is set by the extent of separation of scale between the two classes of reactions. We call the location of the separation the transition line (orange line in Fig. 3B), see section S6.3.5). At large aaRS concentrations, the transition line plateaus as a result of the finite total tRNA budget within the cell (Fig. 3B, middle panel). The plateau is reached once all tRNAs aaRS are charged: the system is then no longer limited by synthetases, but by the amount of tRNAs. In other words, at large aaRS and EF-Tu abundances, the total tRNA concentration becomes limiting for the TC concentration.

The transition line corresponds to conditions in which EF-Tu and aaRS are co-limiting for TC concentration. In the EF-Tu limited region, increasing aaRS abundance does not increase ternary complex concentration: since all EF-Tu proteins are already bound to charged tRNAs, increasing tRNA charging cannot further increase TC concentration. Conversely, in the aatRNA limited region, increasing EF-Tu abundance does not increase TC concentration: since all charged tRNAs are already bound by EF-Tu, increasing EF-Tu concentration does not alleviate the requirement for more charged tRNAs. Given that the optimality condition requires non-zero increase in ternary complex concentration with increasing factor abundance (equations 5 and 10), the optimal EF-Tu and aaRS abundances must be on the transition line. Away from this line, there is either an unproductive excess of EF-Tu or aaRS.

Which point on the transition line corresponds to the optimum? As an approximation, note that inside the EF-Tu limited region, *ϕ*_*TC*_ ≈ *ϕ*_*Tu*_ (since most EF-Tu proteins are bound by charged tRNAs). Hence, within this region, the optimal abundance for the ternary complex can be solved in the same way as for translation termination factors resulting in the optimum:

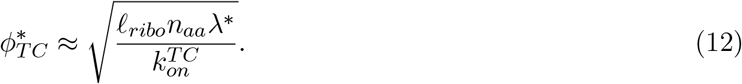

Importantly, compared to the solution for EF-G, the above is multiplied by an additional factor of 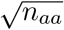. This contribution arises from the rescaling of the association rate for the ternary complex to the ribosome in our coarse-grained one-codon model, reflecting the fact that the interaction between the ternary complex and the ribosome is codon specific.

From the requirement for the combined EF-Tu and aaRS solution to fall on the transition line, the approximate solution for the optimal tRNA synthetase abundance is then the intersection (yellow star in Fig. 3B) of the transition line with the EF-Tu-only solution described above (dashed blue line in Fig. 3B) and is related to the amount of excess tRNA (Supplement, section S6.3.5).

For the above heuristic to be valid, the total number of tRNAs in the cell must be sufficient to accommodate all ribosomes (about 2 per ribosome) and binding to all EF-Tu (about *>* 4 per ribosome based on endogenous expression stoichiometry [34, 38]). The number of tRNAs per ribosomes in the cell should thus be at least 6×. Remarkably, estimates of this ratio in the cell suggest that this is just barely the case (between 6-7 tRNAs/ribosome at fast growth [15]). Although our model treats the total tRNA abundance as a measured parameter and omits its selective pressure (but see [21]), the abundance of three core components of the tRNA cycle appear to be at the special point where the transition line plateau, that is set by total tRNA abundance, just crosses the EF-Tu-only optimum (blue line in Fig. 3B). At this point, all three components are co-limiting.

### Optimal stoichioimetry of mRNA translation factors

Analogous to the case studies above, optimal concentrations for all core translation factors can be solved using equation 3 and their respective kinetics schemes (the case of translation initiation is solved in Supplement, section S7). The analytical forms of the optimal solutions are shown in Table S3. In the diffusion limited regime, the ratios of growth-optimized TF concentrations are independent of the growth rate (except for aaRS), and are dependent only on basic biophysical parameters, such as protein sizes and diffusion constants.

To obtain the numerical values for the optimal TF stoichiometry, we note that the ratios of diffusion-limited biomolecular association rate constants between two reactions can be expressed using Smoluchowski’s formulation as 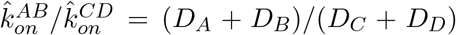 (where the ^ refers to units µM^−1^s^−1^, see Supplement, section S1.2), where *D*_*i*_ is the diffusion constant for the molecular species *i*. Diffusion constants for several TFs have been measured experimentally [4, 55, 59, 67], and other uncharacterized ones can be estimated using the cubic-root scaling with protein size from the Stokes-Einstein relation [49]. Importantly, the absolute values of the optimal concentrations can be anchored by the measured association rate constant between TC and the ribosome obtained from translation elongation kinetic measurements *in vivo* [13] (Supplement, section S9). The square-root dependence on these parameters (Table S3), as explained in the case of termination factors, makes the numerical values less sensitive to the exact biophysical parameters.

The optimal TF concentrations show concordance with the observed ones, both in terms of the absolute levels and the stoichiometry among TFs (Fig. 4). A hierarchy of expression levels emerges such that the factors involved in elongation are more abundant compared to initiation and termination factors. The separation of these two classes is driven by the scaling factor 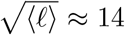 in our analytical solutions, which reflects the fact that the flux for elongation factors is ⟨ ℓ⟩ ≈ 14; times higher than that for initiation and termination factors. Within each class, the finer hierarchy of expression levels can also be further explained by simple parameters. For example, EF-Tu is more abundant than EF-G by a factor of 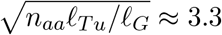 (observed *ϕ*_*Tu*_*/ϕ*_*G*_: *E. coli* 3.9, *B. subtilis* 2.7, *V. natriegens* 3.3). A higher abundance is required for EF-Tu because it is bound to the different tRNAs, which effectively decreases the concentration by a factor of *n*_*aa*_ ≈ 20 (see section S6.2 for derivation and discussion of why the factor is not equal to the number of different tRNAs). Taken together, our model offers straightforward explanations for the observed TF stoichiometry.

**Figure 4:**
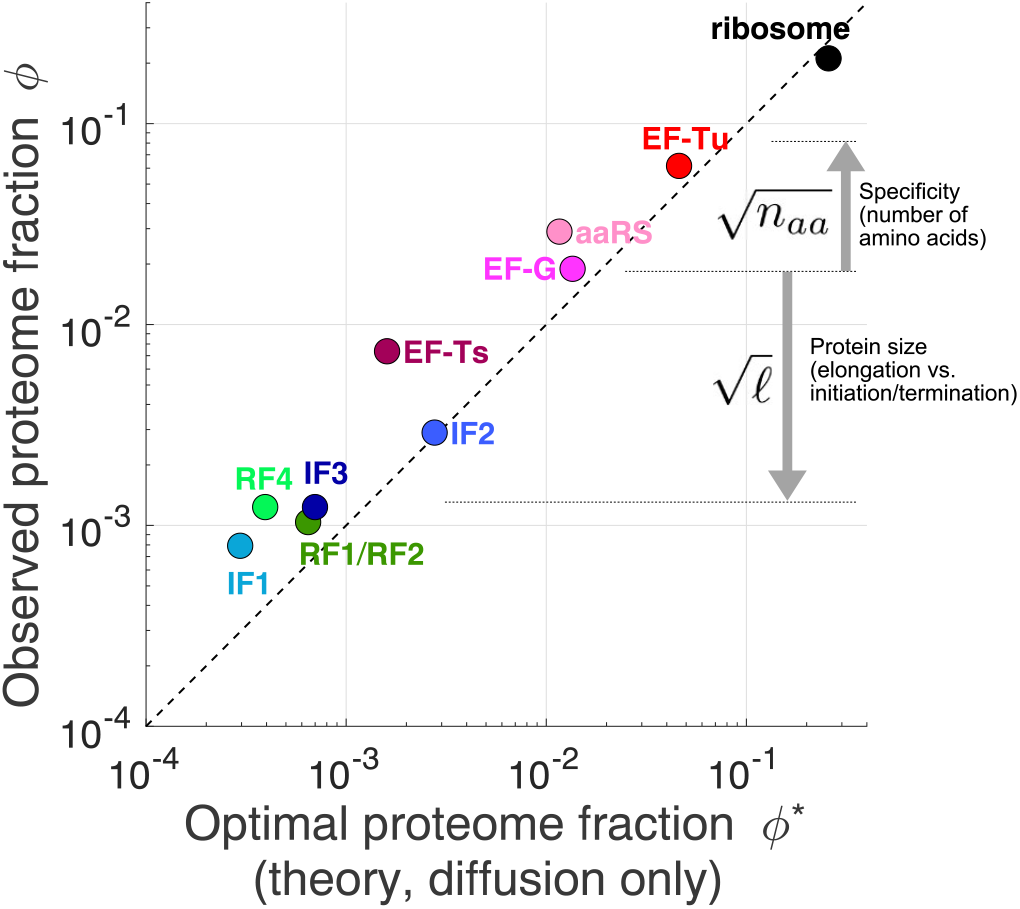
Predicted optimal abundance (no catalytic contribution, *k*_*cat*_ → ∞) versus observed abundance. Experimental values are the average of *E. coli, B. subtilis, V. natriegens* [34]. We note that given the extreme sensitivity of the optimal aaRS abundance on the total tRNA to ribosome ratio (visually: yellow star’s position moves rapidly along x-axis upon changes in plateau of transition line, see Fig. 3B), the agreement for aaRS should be interpreted with caution. Values are listed in Table S5.

For a few TFs, the observed concentrations are 2- to 5-fold higher than the predicted optimal levels (e.g., EF-Ts, RF4, and IF1 in Fig. 4). A potential explanation is that the corresponding reactions may not be diffusion-limited, which would lead to a non-negligible fraction of TFs sequestered at the catalytic step and thereby require higher total concentrations. Our optimization model can also be solved analytically in the non-diffusion-limited regime (Table S3). However, the numerical values for these solutions are difficult to obtain because the estimates for catalytic rates are sparse and often inconsistent with estimates of kinetics in live cells (Supplement, section S9). Another potential explanation is that the selective pressure for these TFs may be lower compared to the more highly expressed TFs. This explanation is unlikely both because their stoichiometry are observed to be conserved (Fig. 1B) and given that the expression of other lowly expressed TFs (e.g., RF1, RF2, and individual aaRSs) has been shown to acutely affect cell growth [33, 52]. Nevertheless, the deviations from the predicted optimal levels suggest that a more refined model may be required than our first-principles derivation.

## Discussion

Despite the comprehensive characterization of their molecular mechanisms, the ‘mixology’ for the protein synthesis machineries inside living cells has remained elusive. Here we establish a first-principles framework to provide analytical solutions for the growth-optimizing concentrations of translation factors. We find reasonable agreements between our parsimonious predictions and the observed TF stoichiometry (Fig. 4). These results provide simple rationales for the hierarchy of expression levels, as well as insights into several construction principles for biological pathways.

An important implication from the agreement between observed stoichiometries and our predictions is that most TFs are co-limiting for growth. Previous models have focused on proteomic allocation constraints to ribosomes [5, 62, 63] and abundant elongation factors [17, 21, 30]. Our results suggest that expression of translation factors, which drive functional outputs from ribosomes, also co-limits cell growth. By virtue of the interlocked translation cycles at steady state, the flux through every cycle must be matched. In our model, the optimality occurs when there are just enough TFs to support the required flux in every cycle, such that the proteome fraction of free factors equals that of waiting ribosomes at that step (equipartition). If the concentration of any one TF falls below the optimal point, it becomes the limiting factor for protein synthesis and growth. This result is supported by experimental evidence that slight knockdowns of individual RFs and aaRSs are detrimental to growth [33, 52]. Figuratively, the translation apparatus is analogous to a vulnerable supply chain, in which slowdown in any of the steps affects the full output.

In the diffusion-limited regime, the optimal TF stoichiometry is independent of the specific growth rate. This is consistent with the observation that relative TF expression remains unchanged between *E. coli* growing with a 20-min doubling time and a 2-hr doubling time [34, 38]. Furthermore, the optimal TF stoichiometry depends only on simple biophysical parameters, including protein sizes and diffusion constants, that are likely conserved in distant species. This result is consistent with the maintenance of the relative TF expression across large phylogenetic distances even though the underlying regulation and cellular physiology has diverged [34]. It remains to be determined if similar biophysical principles apply to the numerous other pathways that also exhibit conserved enzyme expression stoichiometry.

In principle, our model can also make predictions on the growth defects at suboptimal TF concentrations. However, experimentally testing these predictions will be difficult due to secondary effects of gene regulation that are not considered in our model near optimality. For example, we have recently shown that small changes in RF levels lead to idiosyncratic induction of the general stress response in *B. subtilis* due to a single ultrasensitive stop codon [33]. As a result, the growth defect not only arises from reduced translation flux, but is in fact dictated by spurious regulatory connections that are normally not activated when TF expression is at the optimum. We propose that TF expression may be set at the optimal levels as our first-principles model suggests but entrenched by spurious connections in the regulatory network. To predict the full expression-to-fitness landscape away from the optimum, a more comprehensive model may be required to take into account all the molecular interactions in the cell [26, 41].

Our coarse-graining approach has several limitations in its connection to detailed biochemical parameters. First, some of the coarse-grained rate constants remain difficult to estimate, possibly neglecting important features. For example, we do not explicitly consider off-rates in our model. Instead, our parameters correspond to effective rate constants that account for possible sequential binding and unbinding events, i.e., 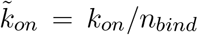, with n_*bind*_ = *k*_*cat*_ */*(*k*_*cat*_ + *k*_*off*_). The effective association rate constants in our model thus contain information about catalytic and possible proofreading steps. We have used a rescaling of *in vivo* measured *k*_*on*_ as an attempt to circumvent this issue, but the specifics of off and/or catalytic rates for each factor could complicate this rescaling in a way that is difficult to predict. Second, the model does not account for possible sequestration of binding sites by non-cognate interactions and it implicitly assumes the off-rates to be large in these cases such that a cognate binding site is never occupied by a non-cognate binding event. An expanded theory adding requirements of accuracy [17, 32, 40] and its interplay with the framework defined here, would be an interesting future research avenue.

Taken together, our model provides the biophysical basis for the stoichiometry of translation factors in living cells. The first-principles approach complements more comprehensive models that include many biochemical parameters [21, 66], while providing intuitive rationales for the expression hierarchy. We anticipate that our approach will be generalizable to elucidate or design enzyme stoichiometry of other biological pathways, especially those whose activities are required for cell growth.

## Acknowledgements

We thank R. Battaglia, J. Cascino, M. Gill, M. Parker, D. Parker, and G. Schmidt for critical reading of the manuscript, and all members of the Li lab for discussion. This research was supported by NIH grant R35GM124732, the NSF CAREER Award, the Smith Odyssey Award, the Pew Biomedical Scholars Program, a Sloan Research Fellowship, the Searle Scholars Program, the Smith Family Award for Excellence in Biomedical Research; NSERC doctoral Fellowship and HHMI International Student Research Fellowship (to J.-B.L.).

## Data availability

Computer scripts used in this study (custom Matlab scripts) will be made available upon reasonable request. Already publicly available ribosome profiling datasets were used (GEO accessions GSE95211 and GSE53767).

## S1 Methods details

### S1.1 Average protein size ⟨*ℓ*⟩

We calculate the average protein size weighted by expression as ⟨*ℓ*⟩ := (Σ _*i*_ e_*i*_)^−1^ *e*_*i*_*ℓ*_*i*_, where *ℓ*_*i*_ is the number of amino acid for the protein product of gene *i*, and *e*_*i*_ is the protein synthesis rate (as estimated from ribosome profiling [34, 38]) for gene *i*. For a stable proteome (in fast growing bacteria, the cell doubling time is shorter than the active degradation of most proteins [35]), the protein synthesis rate equals to the proteome mass fraction [38]. Changes in the expression of genes across growth conditions do not lead to substantial changes in ⟨*ℓ*⟩. In *E. coli*, across growth conditions spanning ≈ 20 min doubling time to ≈ 120 min, ⟨*ℓ*⟩ changes by about 20%. Specifically, we find ⟨*ℓ*⟩ =196 a.a., 210 a.a., and 240 a.a. in respectively MOPS complete (≈ 20 min doubling time [38]), MOPS minimal (≈ 56 min doubling time [38]), and NQ1390 forced glucose limitation (≈ 120 min doubling time, unpublished), based on ribosome profiling data.

### S1.2 Conversion between concentration and proteome fraction

Throughout, we use both units of concentration (molar), denoted as e.g., [*A*] for protein *A*, and proteome fraction, denoted by *ϕ*_*A*_ [62]. The correspondence between the two is *ϕ*_*A*_ = [*A*]*ℓ*_*A*_*/P*, where *ℓ*_*A*_ is the number of amino acid in protein *A*, and *P* is the in-protein amino acid concentration in the cell. *P* ≈ 2.5 × 10^6^ µM, and has a value approximately independent of growth rate [11, 30]. This change in units also relates to how association constants are defined in units of proteome fraction: 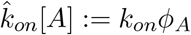, where the hat 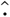 refers to the association constant in usual units of µM^−1^ s^−1^ (used to connect to empirical data). Hence, 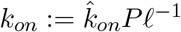 is the rescaled association rate in units of proteome fraction.

### S2 Equality of ribosome flux in steady-state

In steady-state exponential growth, the ribosome flux in and out of each intermediate state is equal to the total flux. This results from the fact that no ribosome can accumulate in any intermediate state. Since the flux out of state *i* is given by 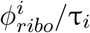, we must have:

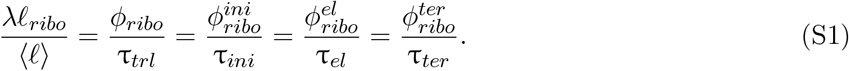

As a consequence, the proportion of ribosome in each state is equal to the proportion of time spent at that given step, for example for translation initiation:

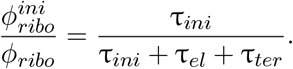

### S3 Coarse-grained transition times: models of ribosome traffic

Our coarse-grained model of ribosome transitions between categories of initiation, elongation, and termination need to be distinguished from the individual molecular times of the respective steps in one important regard: ribosome traffic on mRNAs can lead to effective delays arising from transient queuing. For example, if translation termination is slow and ribosomes start to pile up and form queues upstream of stop codons on mRNAs, the molecular time of termination (time between ribosome arrival to the stop codon and its recycling to the free ribosome pool) will not be a correct reflection of the actual termination time of a ribosome, because of the additional wait time in the queue. A similar argument can be made for transient queuing forming in the body of genes for elongating ribosomes.

We connect these two (molecular and coarse-grained) levels of description by noting that our mass action schemes relating the translation factor abundance to the times of the specific steps can be used as input parameters in traffic models of ribosome movement along mRNAs taking into account possible many-body interactions (e.g., totally asymmetric exclusion processes [27, 64]). Solving these traffic models can then be used to obtain transition times in our coarse-grained translation cycle model. As we show below, corrections arising from transient queuing are small (for endogenous translation factor abundances) based on current estimates the absolute rates of initiation, elongation, and termination, on individual mRNAs, such that stochastic queuing does not play a dominant role in determining optimal translation factor expression levels.

As a first example, we relate the on-stop codon molecular termination time τ_*ter*_, which we obtain from solving our mass action scheme (see equation 6), to the termination time in presence of queuing: 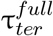. The difference between the two, as described above, being related to possible queues upstream of stop codons leading to further delays in the process of translation termination, and thus to a longer termination time than that of the molecular on-stop codon termination. The delay factor will be denoted 𝒬 (τ_*ter*_), defined through:

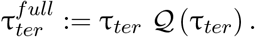

To derive the expression for the 𝒬 factor, note that in steady-state, ribosome numbers in a given state is directly proportional to the time to transition out of that state. Let *m*_*i*_ be the mRNA concentration for gene *i* in the cell, *n*_*ter*_ (*α*_*i*_, τ_*ter*_) the number of terminating ribosomes (including queues if present) on a transcript with per mRNA translation initiation rate (i.e., translation efficiency [37]) *α*_*i*_, then:

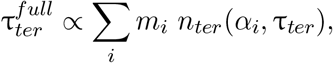

whereas

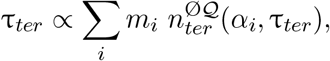

with 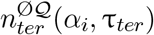 the average number of terminating ribosomes on a transcript with translation efficiency *α*_*i*_, assuming no queue upstream of the stop codon. Note that 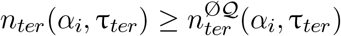 (the differences being queued ribosomes). Hence, the queuing factor 𝒬 is:

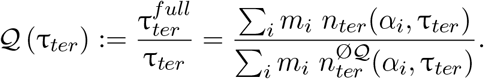

Formally, *n*_*ter*_ can be obtained by solving a TASEP model [64], but a simplified queue model [7, 33] disregarding spatial information recapitulates the statistics of queue formation (as verified by full stochastic simulations, data not shown). The state space of the queue model is the number of ribosomes *N* in the queue. Ribosomes arrive at a rate *α* (initiation rate on the transcript), and leave at the molecular termination rate 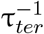. The ribosome arrival rate at the queue is rigorously correct in steady-state, unless the queue becomes large enough to affect the initiation process (fully jammed transcript), or RNA degradation. The stochastic process (away from the jammed state) is then described by: *N* → *N* + 1 at rate *α*, and *N* → *N* − 1 at rate 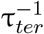 for *N >* 0. The probability for the queue to have *N* ribosomes, *P* (*N*), can be obtained as the steady-state from the resulting master equation, leading to a geometric series: *P* (*N*) = (*α*τ_*ter*_)^*N*^ (1 − *α*τ_*ter*_). Hence, the prevalence of higher order queues scales as the ratio of the initiation to termination rate on the transcript. The average queue size, corresponding to *n*_*ter*_ (*α*_*i*_, τ_*ter*_), is:

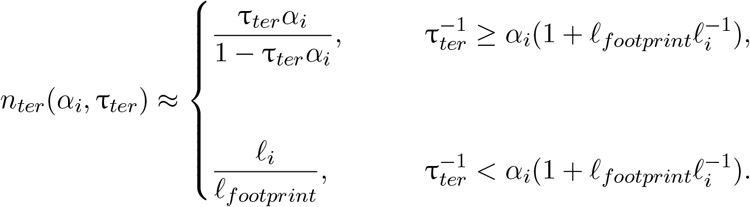

Above, the solution of the simple model is truncated at the value where the transcript becomes fully jammed with *ℓ*_*i*_*/ℓ*_*footprint*_ ribosomes (*ℓ*_*i*_ and *ℓ*_*footprint*_ being the size of gene *i* and the size occupied by a ribosome respectively). The no queue ribosome number is simply equal to a model where queues with *N >* 1 do not arise, hence 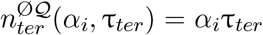. Therefore, the queuing factor, under the stated assumptions (and assuming no transcript is in the jammed state), is

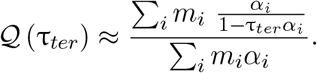

Expanding for fast termination gives 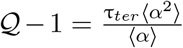 as the leading order correction, where the averages are weighted by mRNA levels. The above was derived assuming exponentially distributed initiation and termination times, but could be modified to account for more complex dynamics of the initiation and initiation steps.

The queuing factor can be estimated based on absolute measurements of the initiation and termination rates in cells. Kennell and Riezman [28] estimate 3.2 s between initiation events on the *lacZ* mRNA (at 48 min per cell doubling). Bremer and Dennis [11] estimate 1 s per ribosome initiation events at 20 min doubling time. Recent calibrated high-throughput measurements report a genome-wide median of 5.6 s per initiation events [19]. To our knowledge, estimation of absolute *in vivo* termination rates have not been performed, but we can estimate bounds. Indirect assessment based on steady-state protein production measurements place the fraction of actively elongating ribosome at about 95% [13]. Assuming (upper bound) that the 5% of non elongating ribosomes are in the process of termination would give a termination time of 5% × 11.1*s* ≈ 0.6 s (fraction of ribosomes in a given state equal to the ratio of transition times), where we have used that the elongation time of an average protein is about 11.1 s (200 *a*.*a*.*/*18 *a*.*a. s*^−1^) at fast growth [13]. This upper bound is still much smaller than the reported median initiation time, suggesting that the queuing factor for termination is small. As additional support to the view that translation is far from being termination limited, small that queues at stop codons are only globally observed in ribosome profiling upon severe perturbations [3, 33, 42, 58]

With regards to translation elongation, transient queuing in the body of gene can also lead to a difference between molecular and coarse-grained transition times in our model. However, the fraction of ribosomes transiently stalled due to this queuing scales as *α*τ_*aa*_ in the low density phase (defined by requirements *α*τ_*ter*_ *<* 1 and 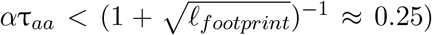 of the TASEP model [64]. Since measured estimates place *α*τ_*aa*_ *∼* 0.01 [13, 19], we do not consider the queuing effect for elongating ribosomes within our optimization framework for elongation factor abundances.

### S4 Protein production flux and growth rate

In order to write the mass action kinetic scheme for more complex models, it is useful to recast our framework in terms of the protein number production flux *J*, defined as the number of full length proteins produced per cell volume per unit time. The production of each protein requires a ribosome to go through the full synthesis cycle, and as such *J* provides a convenient quantity in mass action schemes formulated in molar units.

In steady-state of exponential growth [13, 47, 62], there is a direct relationship between the growth rate *λ* (defined through d*N/*dt = *λN*, where *N* is the number of cells per unit volume of culture) and the protein production flux *J*. Explicitly, the protein mass accumulation rate is *λM*, where *M* is the total protein mass per unit volume of culture. If *V* is the mean cell volume, then *λM/V* = *N m*_*aa*_⟨*ℓ*⟩*J*, where *m*_*aa*_ is the mean amino acid mass. Defining *P* := *M/*(*m*_*aa*_*N V*), the in-protein amino acid concentration per cell (see section S1.2), the connection between protein production flux *J* and growth rate *λ* is then 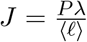. This relationship will be used to convert between molar and proteome fraction in some equations below.

### S5 Translation termination

#### S5.1 Omitted molecular details

The kinetic scheme presented in Fig. 2A does not include some known molecular details of translation termination. For example, GTPase RF3 has been shown to catalyze the release of RF1/RF2 post peptide hydrolysis and to effectively prevent rebinding to empty A site ribosome without peptide [53]. RF3 is not included in our model given our desire for a parsimonious description and due to the absence of identifiable homologs in multiple bacteria (e.g., *B. subtilis*) [43]. Our scheme aggregates the RF1/RF2 recycling rate with the catalytic rate, and further assume a unidirectional reaction without rebinding (consistent with a lower bound), effectively taking into account the action of RF3. In addition, translocation factor EF-G is known to be implicated in ribosome recycling via translocation post RF4 binding [71]. We assume EF-G’s abundance requirement towards the function of termination to be a minor fraction of its total requirement (non-sense to sense codons ≈0.5%) and to be non-limiting for this step. We thus coarsegrain EF-G’s role in ribosome recycling through an effective catalytic rate for RF4, see [10] for details of EF-G’s involvement in ribosome recycling. As another example of simplification in our coarse-graining, we also do not explicitly model RF1/RF2’s post-translational modification by methyltransferase PrmC [48]. Thus, the activity of the RFs within our description to correspond to the average within a possibly heterogeneous pool of modified and unmodified factors in the cell.

#### S5.2 Non-diffusion limited regime (one stop codon)

If translation termination is not diffusion limited, terms corresponding to the finite catalytic times must be included in addition to the diffusive contributions in the termination time (equation 6). Under our simplified scheme (Fig. 2A) and with a single stop codons (grouping RF1 and RF2), the molecular termination time is then sum of the four separate times corresponding to distinct events:

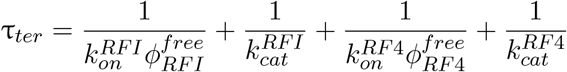

The two novelties compared to the diffusion-limited regime (equation 6) are: (1) addition of the catalytic times 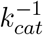 for the two steps, and importantly (2) the mass action diffusion terms now involve the free concentration of release factors. Generally, the free concentration of the TFs can be obtained by solving the steady-state solutions of kinetic schemes under constraints imposed by conservation equations. The examples in e.g., sections S5.3, S6.3, and S7.1 below provide the mathematical details associated with the procedure.

Here, the difference between the total and free concentration of release factor arises from the finite catalytic turnover of the enzymes, and corresponds to the concentration of ribosome bound release factors. Given the flux *J* through the system in steady-state of growth, the concentration of ribosome bound release factor (e.g., for RF4) is 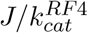, which becomes 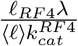 upon converting to proteome fraction. This quantity sets the absolute minimum for the release factor abundance necessary to sustain growth *λ* for a given *k*_*cat*_. The free concentrations for the release factors are then:

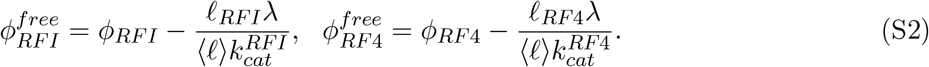

Hence, the final solution for the steady-state termination time as a function of the total abundance of the release factors and growth rate is:

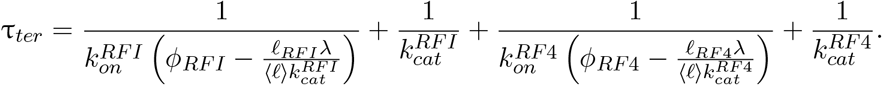

The relationship above, between termination time, total TF abundance, and growth rate *λ* closes the solution of the kinetic scheme. Substituting the above in the optimality condition (equation 5) leads to the solution:

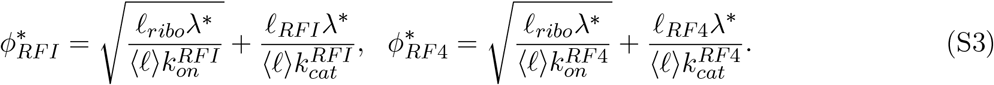

The additional terms ∝ *λ*^*^ correspond to the contribution to the optimal abundance arising from the finite catalytic rates, no present in the diffusion limited regime (equation 7).

#### S5.3 Full three stop codons model

The full model with three different stop codons (UAA, UGA, UAG) and RF1/RF2 with different specificities (RF1: UAA, UAG; RF2: UAA, UGA) can also be solved exactly, leading to a small correction on the summed optimal abundance for RF1 and RF2 of 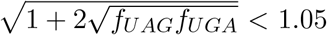 (fast growing species considered, where *f*_*UAG*_ and *f*_*UGA*_ are the fractional fluxes through the RF1 and RF2 stop codons respectively) compared to the single stop codon optimum derived above (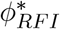, equation S3). We provide details below. With three stop codons, the coarse-grained reaction scheme is shown in Fig. S1. The relevant chemical species and parameters are listed in Table S1.

**Figure S1:**
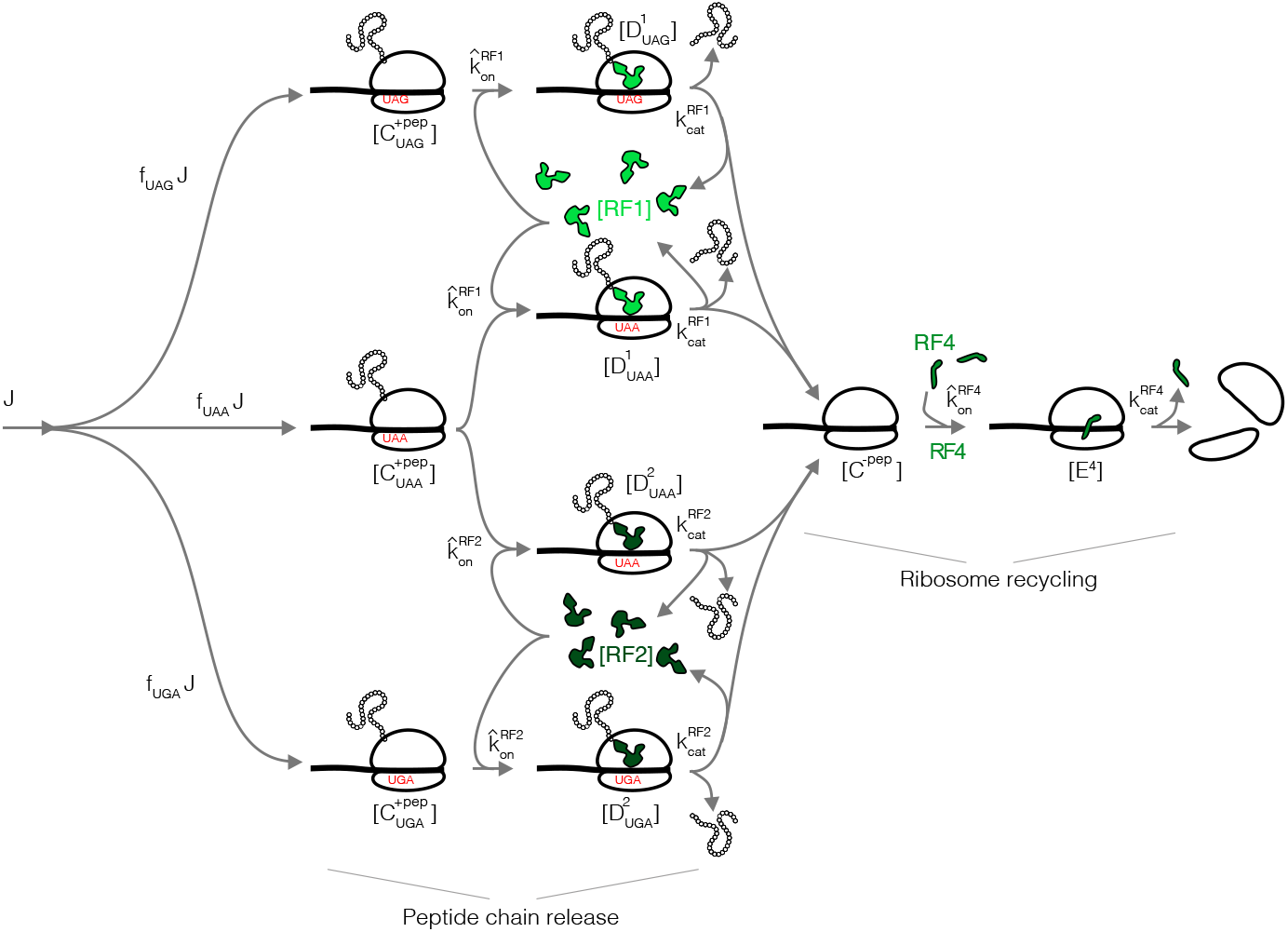
Coarse-grained translation termination scheme with three stop codons and RF1/RF2.

**Table S1:** Chemical species and parameters in three stop codons termination model.

| Variable | Description |
| --- | --- |
| $[C_{UAA}^{+pep}]$ | Ribosomes at UAA with peptide chain $[\mu M]$ |
| $[C_{UAG}^{+pep}]$ | Ribosomes at UAG with peptide chain $[\mu M]$ |
| $[C_{UGA}^{+pep}]$ | Ribosomes at UGA with peptide chain $[\mu M]$ |
| $[D_{UAA}^1]$ | Ribosomes at UAA with peptide chain and RF1 bound $[\mu M]$ |
| $[D_{UAG}^1]$ | Ribosomes at UAG with peptide chain and RF1 bound $[\mu M]$ |
| $[D_{UAA}^2]$ | Ribosomes at UAA with peptide chain and RF2 bound $[\mu M]$ |
| $[D_{UGA}^2]$ | Ribosomes at UGA with peptide chain and RF2 bound $[\mu M]$ |
| $[C^{-pep}]$ | Ribosomes at all stops without peptide chain $[\mu M]$ |
| $[E^4]$ | Ribosomes at all stops without peptide chain and RF4 bound $[\mu M]$ |
| $[RF1]$ | Free RF1 $[\mu M]$ |
| $[RF2]$ | Free RF2 $[\mu M]$ |
| $[RF4]$ | Free RF4 $[\mu M]$ |
| $J_{UAA} = f_{UAA}J$ | Ribosome flux through UAA $[\mu M s^{-1}]$ |
| $J_{UAG} = f_{UAG}J$ | Ribosome flux through UAG $[\mu M s^{-1}]$ |
| $J_{UGA} = f_{UGA}J$ | Ribosome flux through UGA $[\mu M s^{-1}]$ |
| $\hat{k}_{on}^{RF1}$ | On-rate for RF1 $[\mu M^{-1} s^{-1}]$ |
| $\hat{k}_{on}^{RF2}$ | On-rate for RF2 $[\mu M^{-1} s^{-1}]$ |
| $\hat{k}_{on}^{RF4}$ | On-rate for RF4 $[\mu M^{-1} s^{-1}]$ |
| $k_{cat}^{RF1}$ | Catalytic rate for RF1 $[s^{-1}]$ |
| $k_{cat}^{RF2}$ | Catalytic rate for RF2 $[s^{-1}]$ |
| $k_{cat}^{RF4}$ | Catalytic rate for RF4 $[s^{-1}]$ |
| $RF1_{tot}$ | Total RF1 $[\mu M]$ |
| $RF2_{tot}$ | Total RF2 $[\mu M]$ |
| $RF4_{tot}$ | Total RF4 $[\mu M]$ |

The corresponding mass action system of equations for peptide release:

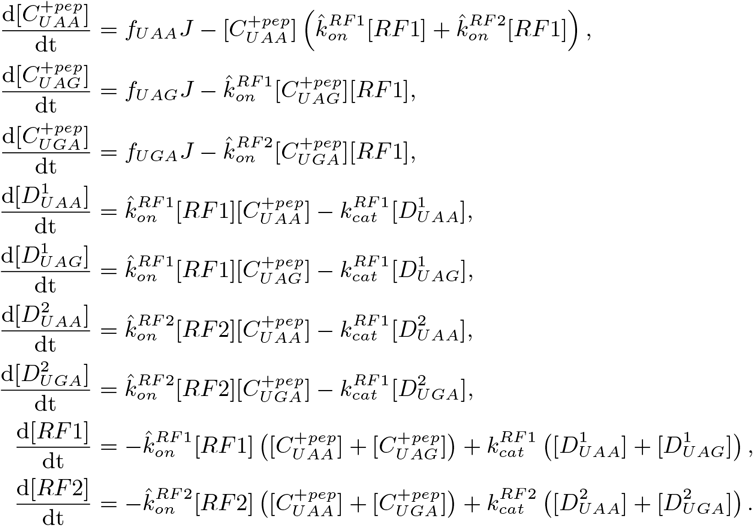

And for ribosome recycling:

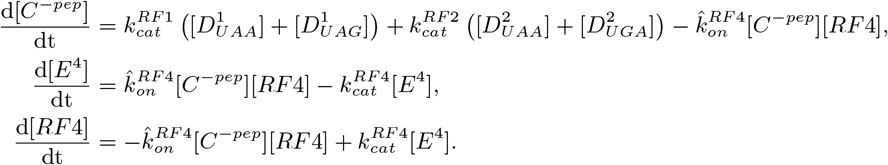

The conservation equations for RF1, RF2 and RF4 are:

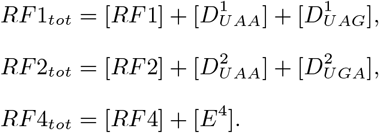

With a more complex scheme such as the one above, the optimization problem can be solved in three steps. First, we obtain the steady-state concentration of the chemical species. Second, we determine the effective coarse-grained termination time. Finally, the optimal abundance is found by substituting the termination time in the optimality condition (equation 5), and solving the resulting system of equation.

##### S5.3.1 Steady-state concentrations for RFs

Note that the RF1/RF2 and RF4 completely decouple, and that the solution for RF4 is identical to the one stop codon case solved above (section S5.2). For peptide chain release, the steady-state of the system can be solved by expressing the all chemical species in terms of [*RF* 1], and [*RF* 2]:

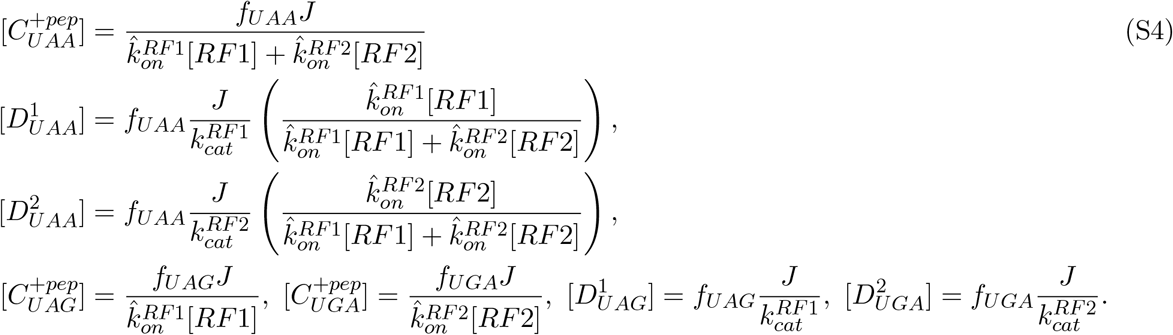

Substituting these in the conservation equations for RF1 and RF2 leads to a closed system in terms of [*RF* 1] and [*RF* 2]:

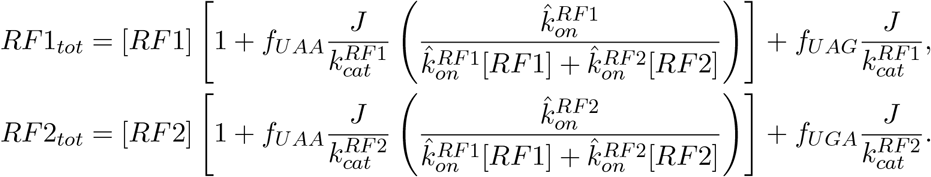

Under the assumption of identical biochemical properties for RF1 and RF2, namely 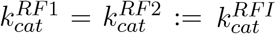 and 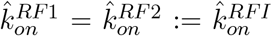, the total free concentration of RF1 and RF2 simplifies to: 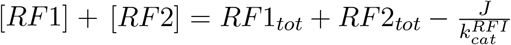, where we used *f*_*UAA*_ + *f*_*UAG*_ + *f*_*UGA*_ = 1 (by definition). Using this relation to eliminate [*RF* 2] from the [*RF* 1] equation (and vice-versa), we obtain, upon conversion to proteome fraction:

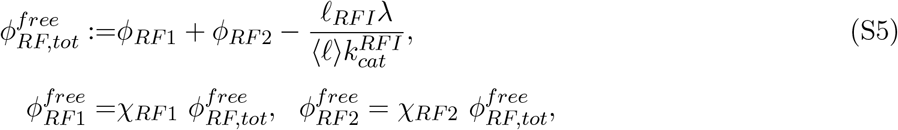

where

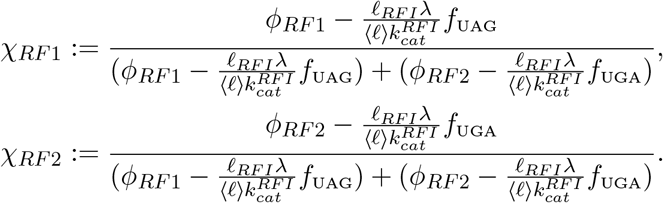

These constitute the steady-state solutions of the system of equation.

##### S5.3.2 Coarse-grained translation termination time

In order to obtain an expression for the termination time (peptide release portion), needed to determine the optimal RF abundance (i.e., to substitute in equation 5), the peptide chain release contribution arises from the ribosome containing species listed in equation S4, which sum to (under the assumption of identical biochemical properties for RF1/RF2):

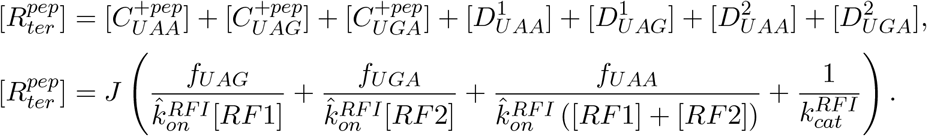

Upon conversion to proteome fraction, the above becomes:

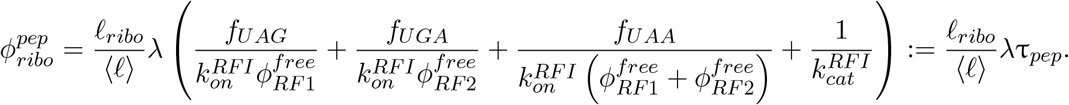

The bracketed term corresponds to the coarse-grained time associated with peptide chain release τ_*pep*_, and the free concentrations are given by equations S5.

##### S5.3.3 Optimal abundances for RF1/RF2

The solved concentrations in steady-state (as a function of proteome fractions) and coarse-grained times allow us to determine the optimal RF1 and RF2 solutions (within our model). The optimality condition (equation 5) is now:

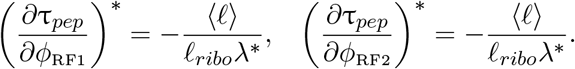

Solving the above system leads to optima 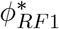 and 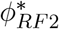:

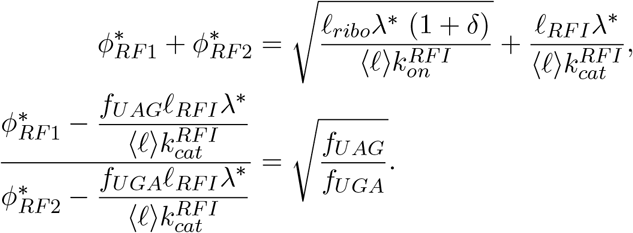

where the new factor 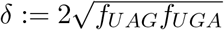.

The flux through each stop codon can be estimated in a variety of bacteria from ribosome profiling data [34] as the total synthesis fraction of genes with the respective stop codon. Results in Table S2 highlight that the three stop codon correction 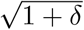 to the optimal solution for the summed abundance of RF1 and RF2 is small for fast growing bacteria considered in the current study.

**Table S2:** Translation flux through stop codon and estimated correction factor 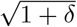, with 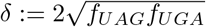.

| Species | $f_{UAA}$ | $f_{UAG}$ | $f_{UGA}$ | $\sqrt{1 + \delta}$ |
| --- | --- | --- | --- | --- |
| *E. coli* | 0.888 | 0.015 | 0.097 | 1.04 |
| *B. subtilis* | 0.888 | 0.064 | 0.049 | 1.05 |
| *V. natriegens* | 0.929 | 0.041 | 0.031 | 1.04 |

### S6 Translation elongation

#### S6.1 Coarse-grained one-codon model

Translation elongation is a more complicated process than termination, involving multiple factors to bring the charged tRNA to the ribosome (EF-Tu), charge the tRNAs (aaRS), translocate the ribosome (EF-G), and perform nucleotide exchange on EF-Tu to drive the process (EF-Ts), in addition to others not included here. Our simplified kinetic scheme is illustrated in Fig. S2. In anticipation coarse-graining procedure detailed below, rates rescaled in the conversion to a one-codon model are marked by *.

To simplify our model, we coarse-grain the elongation cycle by considering a single codon type (section S6.2 below or details of the coarse-graining procedure), effectively grouping the tRNA’s, tRNA synthetases, and different ternary complexes to single entities. Importantly, as a result, the on-rates associated with these processes are rescaled by a factor close to 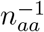, where *n*_*aa*_ = 20.

An important distinction for elongation compared to initiation and termination is that multiple elongation steps (average ⟨*ℓ*⟩ ≈ 200 a.a.) are required to generate a protein. Hence, the flux into the through the elongation cycle is ⟨*ℓ*⟩ larger than that through the initiation and termination steps (there is one initiation and termination event for each protein made, but about 200 elongation steps on average).

**Figure S2:**
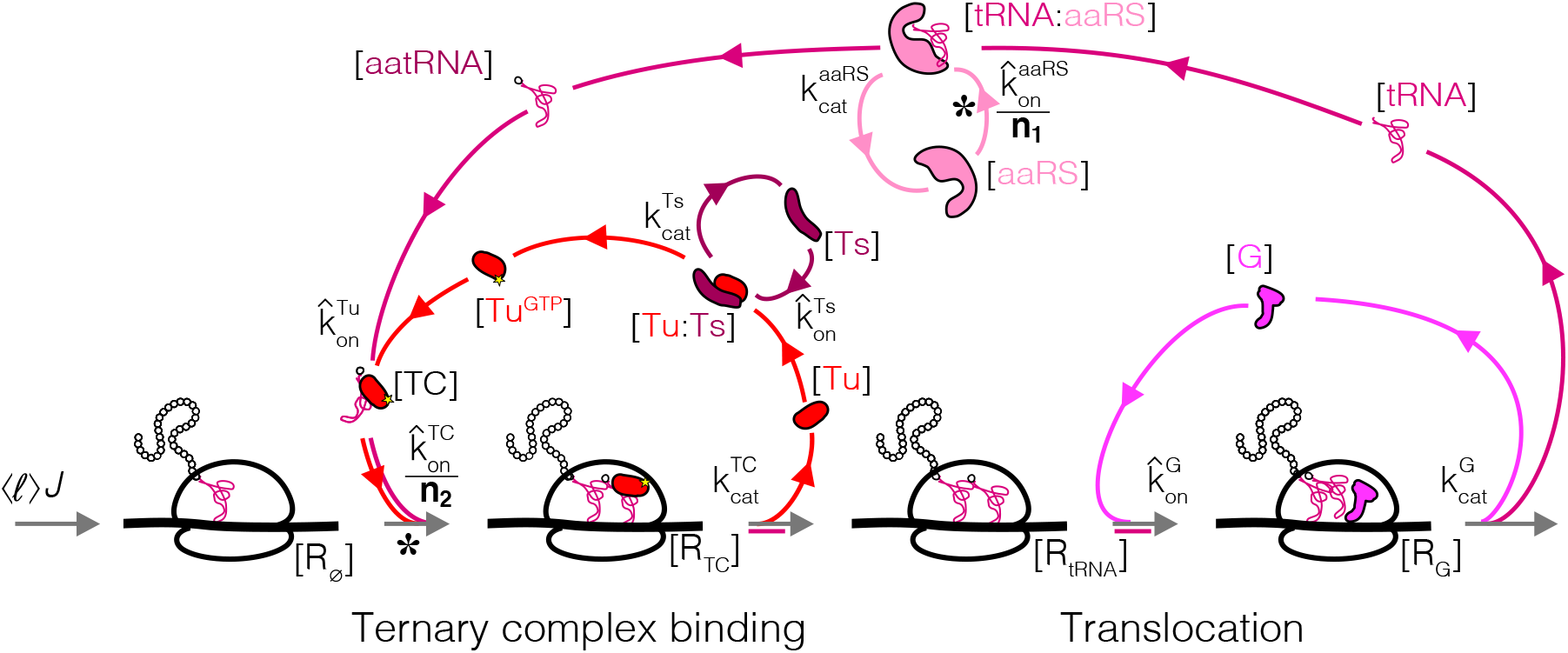
Coarse-grained reaction scheme for a single step (amino acid incorporation) of translation elongation. Tu: EF-Tu, Ts: EF-Ts, G: EF-G, aaRS: aminoacyl tRNA synthetases. Steps with slower rates as a result of the coarse-graining to one effective codon are marked by *.

The mass action reaction scheme for translation elongation:

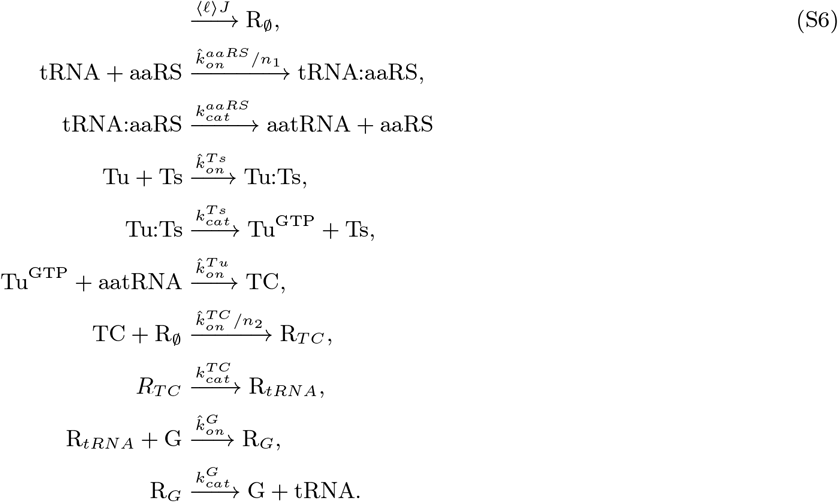

To arrive at the above, we started with a full model of translation (not shown), will all possible codons, tRNA species, and ribosomes with different codons. To coarse-grain the model, we introduced the following effective variables, which correspond to the total concentration of each type of species involved, summed over the of the codon/amino acid specificity:

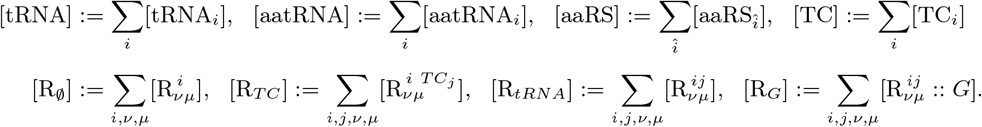

In the above, Greek indices correspond to different codons on mRNAs, and Roman indices to different tRNAs. Roman indices with a hat 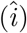 correspond to tRNA synthetases recognizing specific tRNAs (multiple amino acids have more than one tRNA isoacceptor). In defining these coarse-grained species (our approach is analogous to that of [13]), we redefined the two following kinetic parameters:

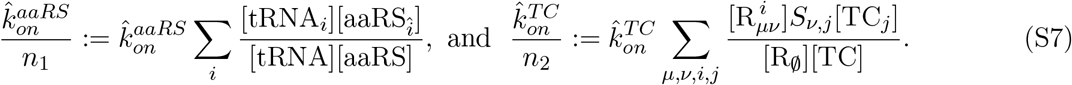

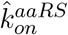 and 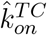 correspond to the microscopic bimolecular rates (assumed equal for the different chemical species). *S*_*v,j*_ is the tRNA isoacceptor/codon specificity matrix (1 if tRNA *i* can recognize codon *v*, 0 otherwise) [9]. Rescaling terms *n*_1_ and *n*_2_ are estimated below.

#### S6.2 Estimation of coarse-grained rates

The definition of coarse-grained parameters (equations S7) involves sums:

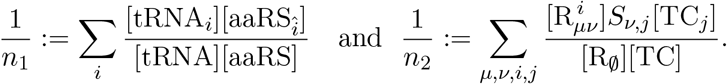

These can be estimated from tRNA abundances, codon usage and individual synthetases’ levels obtained from ribosome profiling data in *E. coli* [38].

We first consider *n*_1_. Note that the fraction of free tRNA of type *i* to the total number of free tRNA (not bound to any protein) is not readily measurable. Assuming similarities between types of tRNA’s, we approximate this fraction with the fraction of total tRNA of type *i* to the total tRNA concentration, or

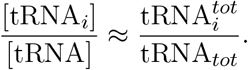

The total tRNA concentration has been measured at fast growth for *E. coli* [15]. The relative concentration of each tRNA synthetases (appropriately corrected for stoichiometry for the different classes) can be computed from the ribosome profiling data [38], and we obtain

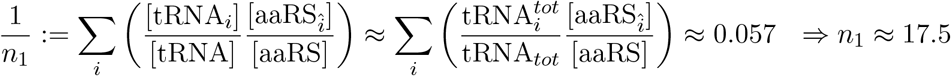

This was to be expected since the synthetases in *E. coli* show little variability around their mean, and in the case of equal synthetase concentration, *n*_1_ = 20 would strictly hold.

For the second sum (*n*_2_), we use distribution of ribosome footprint reads across the transcriptome to estimate ribosome occupancies at different codons. We first make the following approximation for one of the sub-sum:

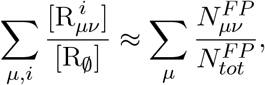

where 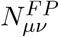 is the total number of ribosome footprint reads at codon pairs *μ, v* and 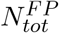 is the total number of footprint reads mapping to coding sequences. The nature of the approximation is that we are taking relative fraction of ribosome footprints (representing ribosomes across the elongation cycle at that codon pair) at a given codon pair to be equal to the relative fraction of ribosomes waiting for the ternary complex to derliver a tRNA to the A site. The modest differences in elongation rates at different codons seen in ribosome profiling data [46] justify this approximation.

From our data (not shown), we have that

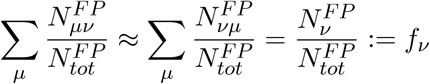

holds to better than 0.5% for each codon. *f*_*v*_ above is the (expression weighted) codon usage. As before with the free tRNA concentrations, we can approximate the relative ternary complexes concentrations by the corresponding total tRNA concentrations:

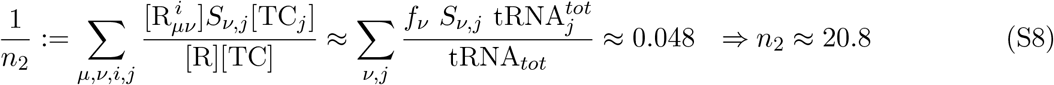

We used the same dataset as before for the total tRNA concentration in *E. coli* [15]. The codon usage was determined directly from ribosome profiling data [38]. The sum of these products is graphically represented in Fig. S3. The above sum of product of tRNA fraction and codon usage provides an effective number of different ternary complexes. *A priori*, that might have been expected to equal to the number of tRNAs (≈40). However, as is apparent in Fig. S3, certain tRNA-codon pairs are much more prevalent than others (even for amino acid with multiple codons and/or tRNA isoacceptors), which leads to a decrease in the effective concentration. The exact value depends on the detailed codon usage and tRNA abundance.

**Figure S3:**
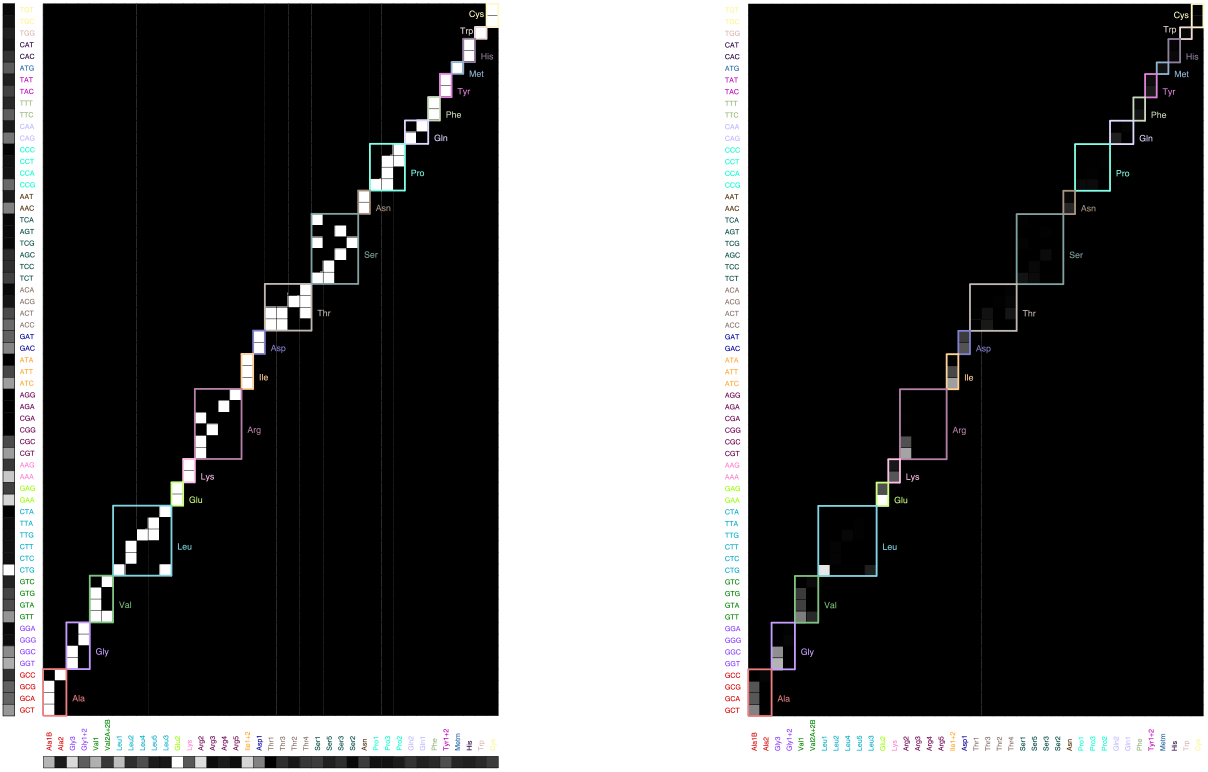
Graphical illustration of the sum (equation S8). Left: codon usage (vertical, from analysis of ribosome profiling data from [38]), tRNA-codon specificity (matrix, from [9], with different amino acids outlined with different colors), and tRNA abundance (horizontal, from [15]) organized by amino acid. Right: product matrix.

Given the results above, we take for simplicity *n*_1_ = *n*_2_ = *n*_*aa*_ = 20.

#### S6.3 Translation elongation: optimal solutions

The mass action reactions corresponding to the one codon elongation cycle model are (equations S6):

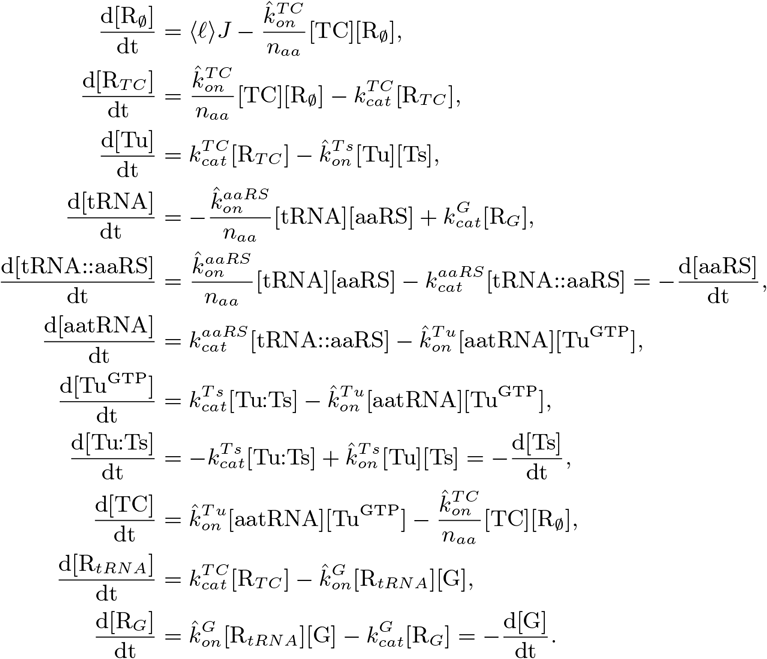

Conservation equations close the system:

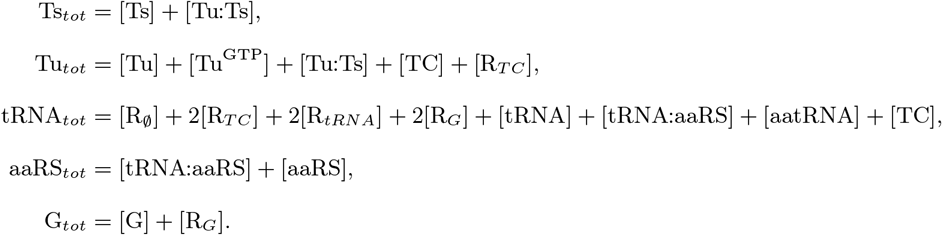

The ternary complex concentration and free EF-G concentration enter the translation elongation time (equation 10, which is the diffusion limited and factor dependent contribution to the elongation time) and are required to infer optimal abundances of elongation factors. Both can to be obtained by solving the system of non-linear equations above.

First, catalytic steps must equal to the flux through in the system in steady-state and thus:

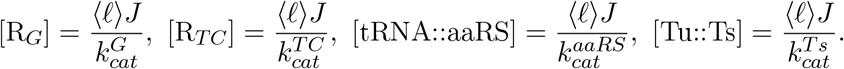

Together with the conservation equations, these allow for immediate solutions for the free concentrations [Ts], [aaRS], and [G]:

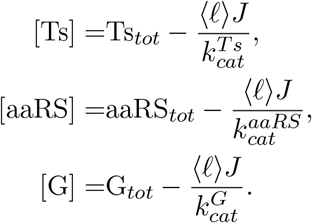

The solution for other species can then also be obtained in terms [Tu^GTP^], and [TC]:

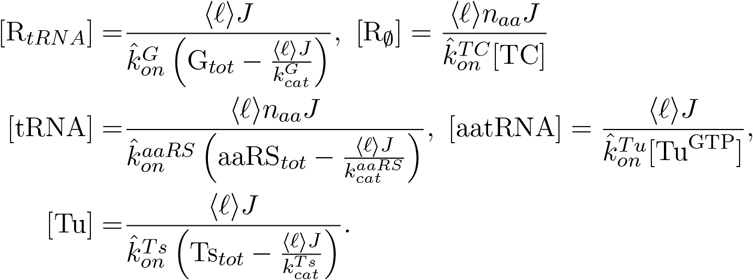

Substituting these in the conservation equations for tRNAs and EF-Tu lead to the final system to solve (converting to proteome fraction):

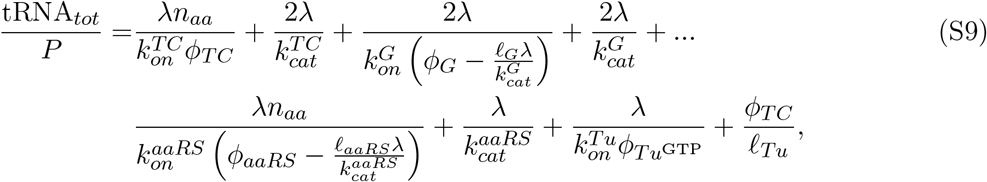

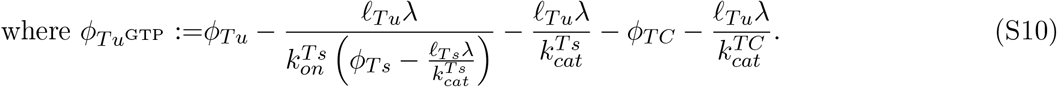

where the solution for 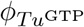 in terms of the ternary concentration was obtained from the conservation equation for EF-Tu. Equations S9 and S10 are closed, and the only variables to solve for is *ϕ*_*TC*_ in terms of the TF abundances: *ϕ*_*Tu*_, *ϕ*_*Ts*_, *ϕ*_*G*_, *ϕ*_*aaRS*_, tRNA abundances, kinetic parameters, and the growth rate *λ*.

#### S6.3.1 Coarse-grained translation elongation time

In order to obtain the coarse-grained translation elongation time, we proceed as for translation termination (section S5.3.2). The summed concentration of the ribosome containing species for translation elongation in our model is:

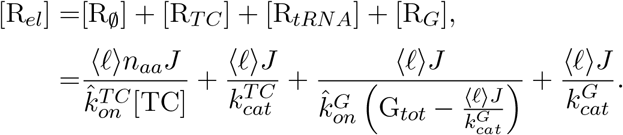

Converting to proteome fraction:

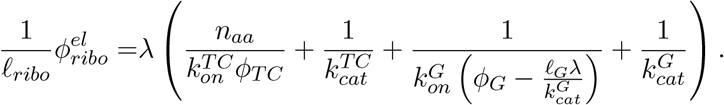

From the coarse-grained flux relations through the different categories (equation S1), which defines the coarse-grained transition times, we thus have:

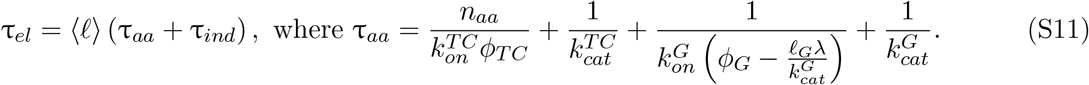

Above, τ_*aa*_ is the effective time for a single step (1 a.a.) of translation elongation, and τ_*ind*_ corresponds to the summed time of factor independent transitions in each elongation step (not explicitly included in the kinetic scheme).

#### S6.3.2 Optimality conditions for translation elongation factors

The optimality condition (equation 5) applied to translation elongation factors leads to:

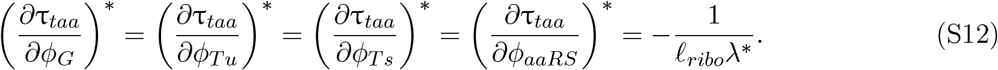

where equation S11 was used for τ_*aa*_. Since the free EF-G concentration does not depend on EF-Tu, EF-Ts, or aaRS concentration, the conditions for EF-Tu, EF-Ts and aaRS simplify to:

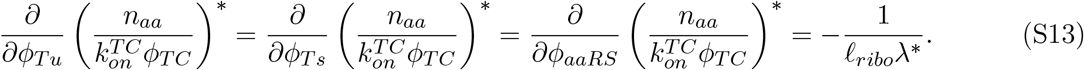

Carrying through the differentiation also leads to conditions on the derivatives of the ternary complex concentration at the optimum:

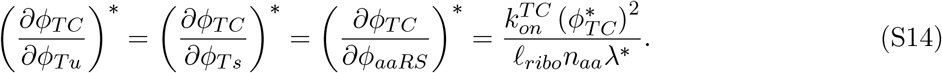

These relationships will be useful to solve for the some elongation factor optimal abundances below.

#### S6.3.3 Optimal EF-Ts abundance

Differentiating equation S9 with respect to *ϕ*_*Tu*_ and *ϕ*_*Ts*_, we get at the optimum:

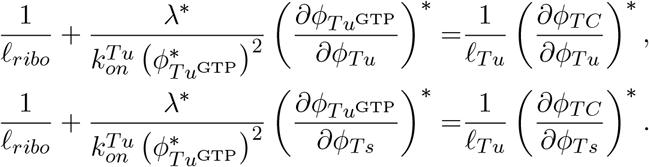

By equation S14, the above leads to the additional condition at the optimum:

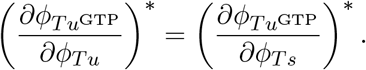

Directly differentiating equation S10, and using equation S14, leads to:

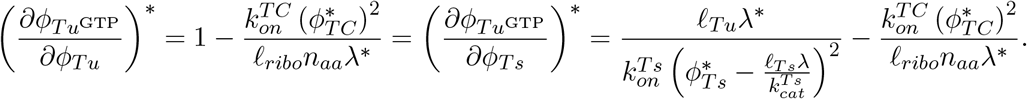

Therefore, the optimal abundance for EF-Ts is:

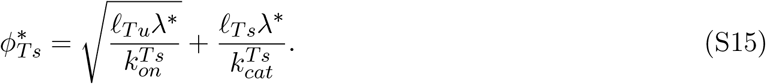

#### S6.3.4 Optimal EF-G abundance

The optimality condition for EF-G is complicated by the fact that EF-G free concentration appears in the solution for the steady-state ternary complex through the tRNA conservation equation S9. Differentiating the conservation tRNA equation, and using the optimality condition S12 (replacing a number of terms with the elongation time τ_*aa*_, equation S11):

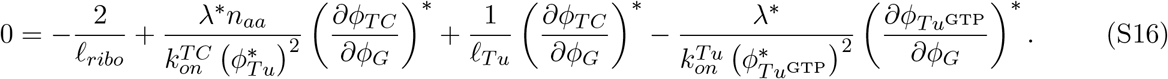

Above, the right-hand portion corresponds to the additional constraint coming from the implication of EF-G in the steady-state concentration of the ternary complex. From the equation for 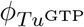 (equation S10), we have directly:

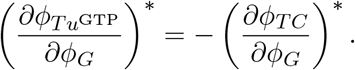

Substituting this in equation S16:

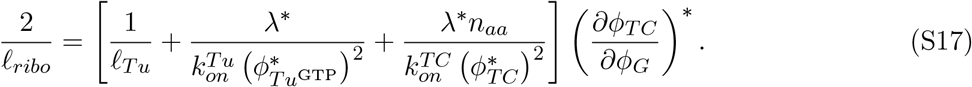

The derivative of the ternary complex with respect to EF-G at the optimum can be obtained from the original optimality condition S12, by carrying through the differentiation:

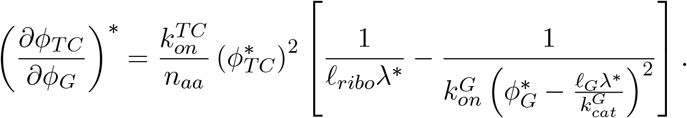

Substituting in equation S17, we arrive at a final equation for EF-G in terms of the concentration of other elongation factor and the optimal growth rate:

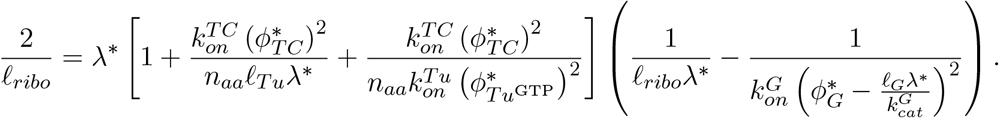

The optimal solution for EF-G is thus:

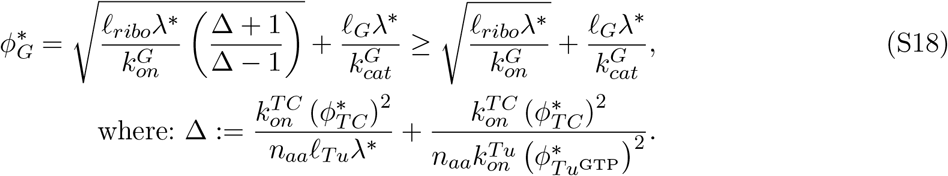

Note that given that the term Δ involves 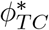 and 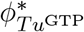, and so the solution above is not a priori complete. However, using the approximate ternary complex concentration at the optimum (equation 12, derived in details in section S6.3.5), we have:

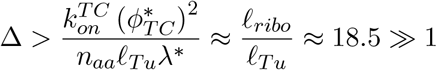

This means that the lower bound for 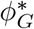 above (equation S18) is a good approximation: in the physiological regime, we can approximately neglect the indirect dependence of the ternary complex concentration on EF-G via the tRNA conservation equation. Hence, the approximate solution for the EF-G optimal abundance is (same for had we initially assumed that *ϕ*_*TC*_ was independent of *ϕ*_*G*_, in which case the solution for EF-G can be obtained identically as that of release factors):

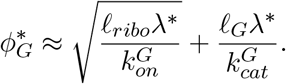

#### S6.3.5 Optimal EF-Tu and aaRS abundances

While simplifying relations were possible with EF-Ts and EF-G, allowing their solution (approximately) independently from the rest of the cycle, EF-Tu and aaRS are intricately connected through the tRNA cycle. We thus return to the tRNA conservation equation, equation S9. For notational simplicity, we group the catalytic step of the TC, EF-G binding, and EF-G catalytic action (translocation) in parameter 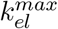 (these do not depend on *ϕ*_*Tu*_ and *ϕ*_*aaRS*_). Further dropping the EF-Ts related and catalytic terms (will be added back at the end, they only contribute a fixed term at the optimum) in the equation for the free EF-Tu, we get:

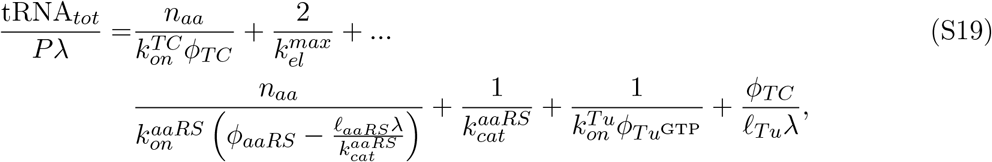

where 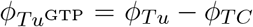 is the free EF-Tu concentration.

This system is first solved numerically (Fig. 3B). To close the equation in terms of uniquely *ϕ*_*TC*_, we use our relationship for *λ* (equation 1), with:

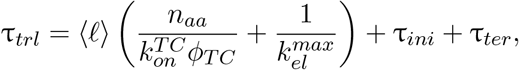

where 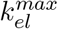 is the maximum rate of translation elongation (contribution other than ternary complex diffusion) estimated from *in vivo* kinetic measurements (≈ 22 s−1 [13]), and τ_*ini*_ + τ_*ter*_ ≈ 0.5 s the estimated time for the initiation and termination step (≈ 5 − 10% of the full translation cycle translation time), taken as fixed parameters here. Using this relationship for the translation time leads to the explicit relationship between growth and ternary complex concentration:

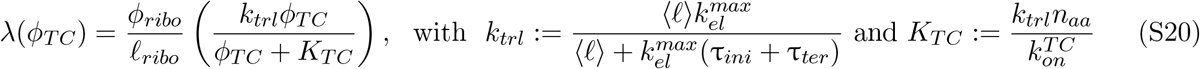

which is the same relationship as the one derived in [30], with the addition of the terms corresponding to the rest translation cycle. Substituting the explicit relationship between growth and ternary complex concentration above (equation S20) in the aaRS/EF-Tu tRNA cycle relationship (equation S19) closes the system for *ϕ*_*TC*_. Numerical solution for this equation is presented in Fig. 3B (see section S9 for other parameters).

The main conclusion from numerically solving the reduced system (equations S19 and S20) is that the EF-Tu/aaRS space is partitioned in two regimes, resulting from the separation of scale of reactions in the coarse-grained model. Specifically, 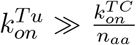, so that any imbalance between the constituents of the ternary complex (charged tRNAs, free EF-Tu), results in stoichiometric unproductive excess of the component in surplus.

We can derive a relation for the “transition line” in the aaRS/EF-Tu space where both free charged tRNAs and free EF-Tu are at low concentrations. This corresponds to setting the (formally impossible) requirement 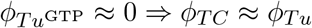 and [aatRNA] 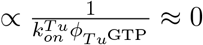, i.e.,

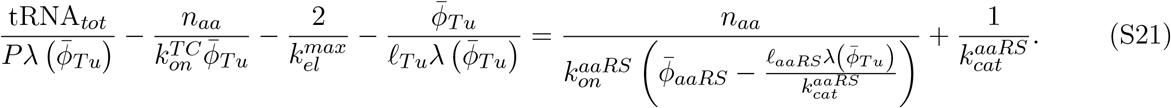

The 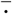 signifies the transition line relationship between 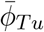 and 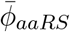, which is displayed in Fig. 3B.

The heuristic to estimate the optimal EF-Tu concentration described in the main text can be extended to include the EF-Ts cycle. In particular, in the EF-Tu limited regime, with 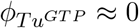, we have (from equation S10):

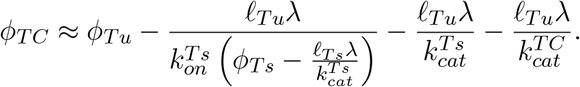

Substituting the above expression for *ϕ*_*T C*_ in the optimality condition (equation S13) for *ϕ*_*Tu*_, we arrive at (using the optimal solution for EF-Ts, equation S15):

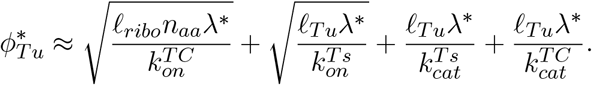

Above, the last three terms (not appearing in equation 12) correspond to the additional diffusion of the EF-Ts cycle, and catalytic contributions.

Following the argument (see main text) that the optimal aaRS abundance should lie on the transition line (equation S21), we obtain:

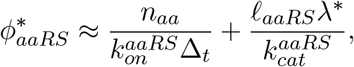

with Δ_*t*_ related to the excess tRNA:

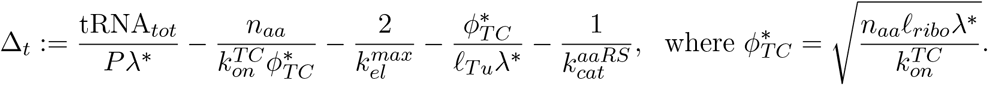

## S7 Translation initiation

Translation initiation is also relatively complex compared to translation termination. In contrast with other steps of the translation cycle, binding of factors necessary for the process (IF1, IF2, IF3, initiator tRNA) do not occur in a strict sequential order, leading to a “heterogeneous assembly landscape” [12, 20] more complex to model. However, one assembly pathway is kinetically favored [45]. We take this favored assembly pathway as our kinetic scheme (Fig. S4, note that binding of tRNA/mRNA are coarse-grained to a single even without loss of generality). We provide some evidence below that taking a more complex assembly pathway would minimally affect the predicted optimal initiation factor abundances.

**Figure S4:**
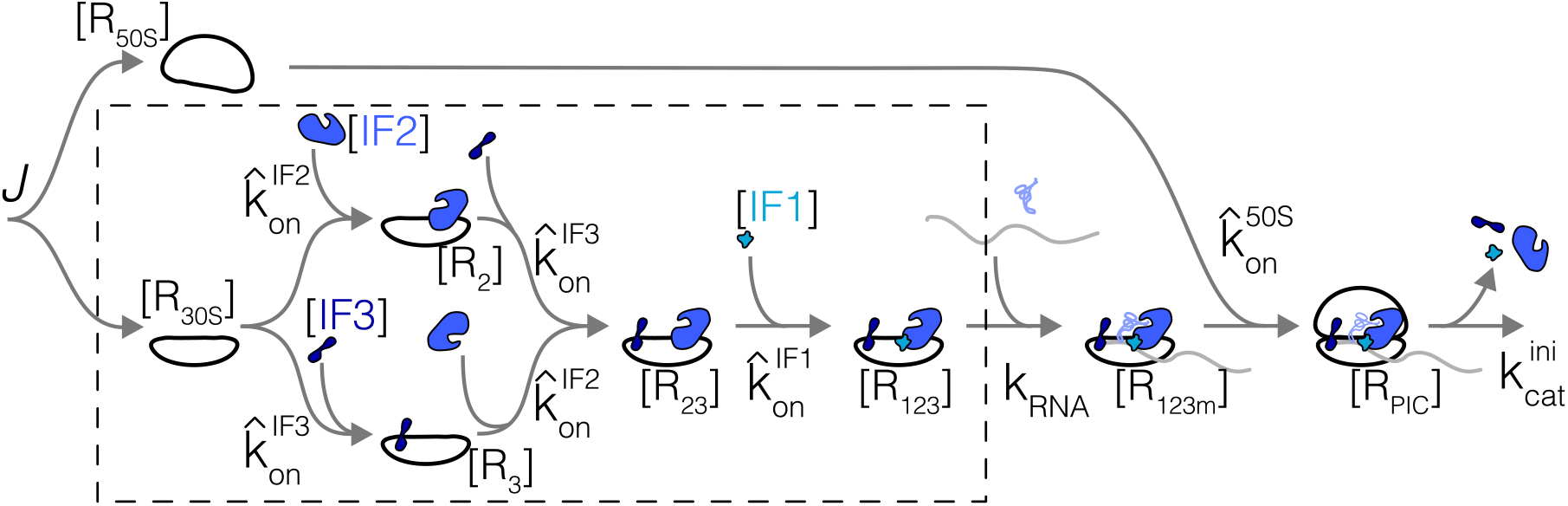
Simplified kinetic scheme for translation initiation. Reactions in dashed box correspond to sub-system solved in detail first (section S7.1). Variables are labeled on the scheme.

The reactions in our simplified schemes are:

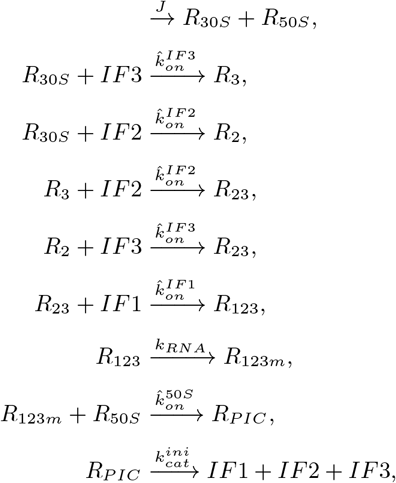

with corresponding mass action equations:

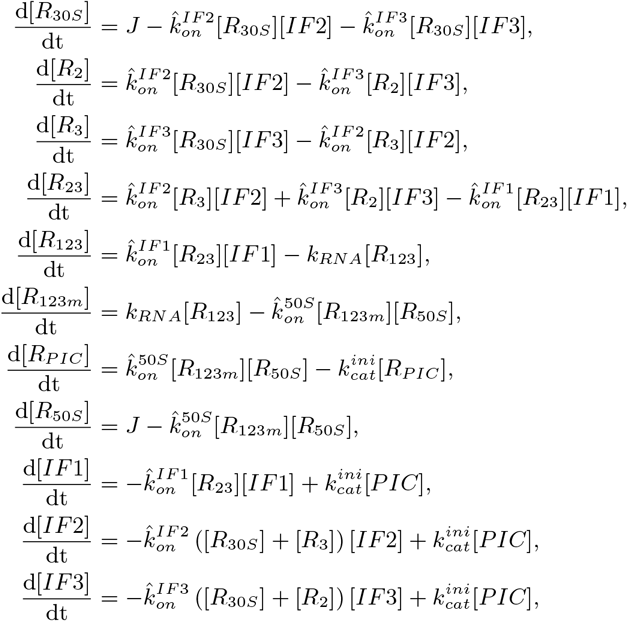

and conservation equations:

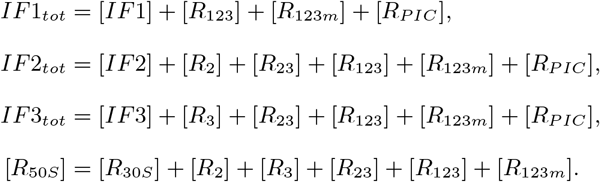

We assume the steady-state concentrations of small and large ribosomal subunits to be equal.

### S7.1 Sub-pathway without subunits joining

The system of equation is complicated by the second branch of the pathway corresponding to 50S subunit binding. However, in the regime 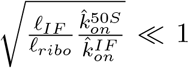 (which is realized because of the large size of the ribosome and slower association rate constant for the large subunit compared to the initiation factors again due to size), the effect of this branch is to add a term to the optimal abundance equal to the concentration of species *R*_123*m*_ (see derivation in section S7.2). We focus here on the solution of the part of the reaction scheme boxed in Fig. S4. This sub-scheme corresponds to:

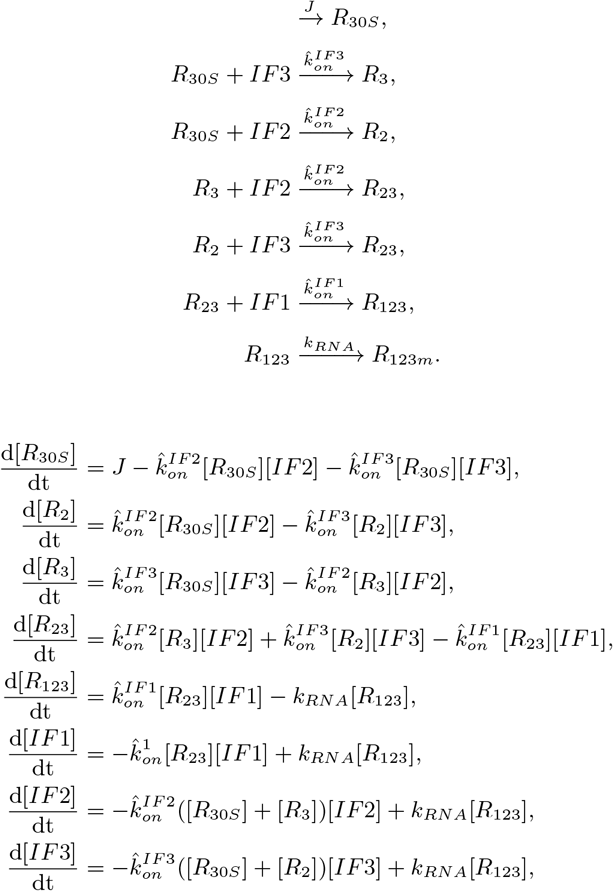

with conservation equations:

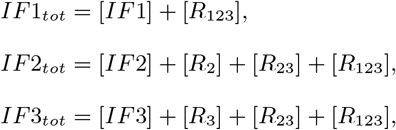

This system can be solved as with the previous schemes. In steady-state, we find for concentrations in terms of the free concentrations [*IF*2] and [*IF*3]:

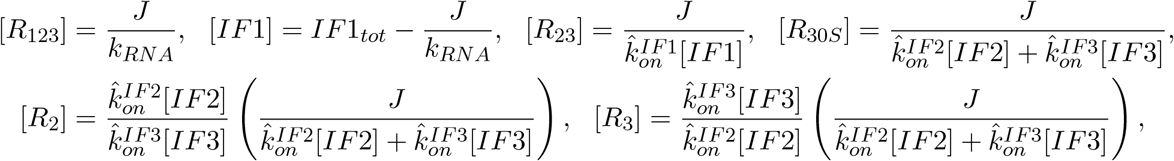

and the coupled equations for [*IF* 2] and [*IF* 3] that need to be solved:

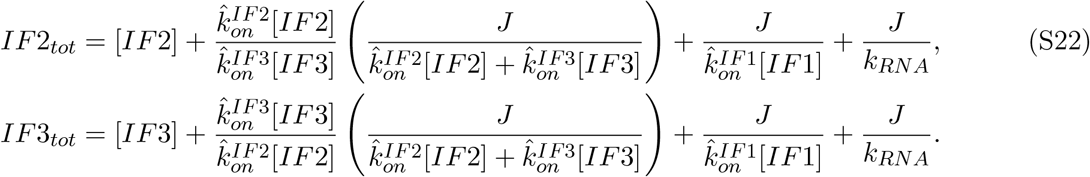

As for translation termination (section S5.3.2) and elongation (section S6.3.1), summing the ribosome containing species:

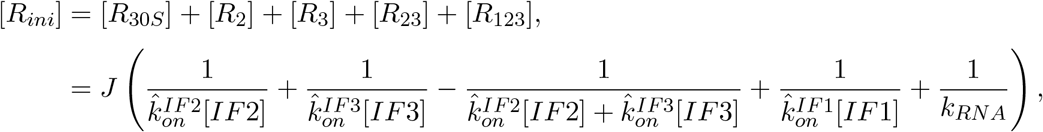

allows us to read the initiation time directly (recast in proteome fraction units):

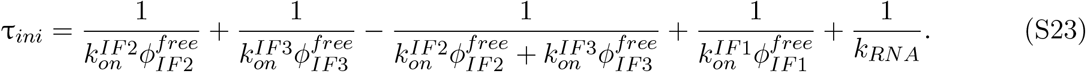

The above is the time can be used in the optimality condition (equation 5). Note that the parallel nature of the reactions with IF2 and IF3 leads to a reduction compared to a purely sequential pathway (negative term above decreasing the total initiation time, as expected if multiple reactions can occur in parallel).

Given that binding of IF1 occurs last in this scheme, its free concentration takes a simple form 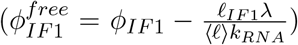. In contrast, computing the free IF2 and IF3 concentrations requires solving the non-linear coupled system, equations S22. Recasting these in units of proteome fraction:

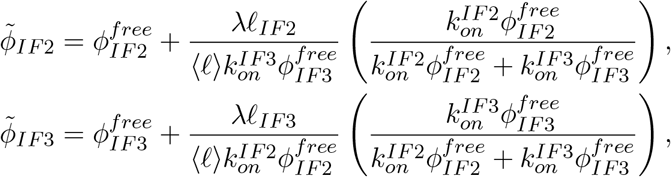

With 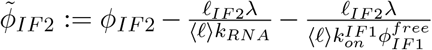, and similarly for 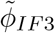. We show now that the terms coupling the two equations for 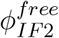 and 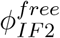 (bracketed above) are small at the optimum. Indeed, based on results in simpler schemes (self-consistency confirmed below), we expect at the optimum:

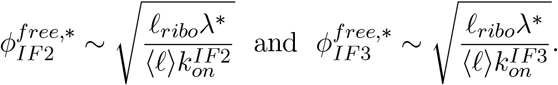

Hence, we expect the two terms at the optimum in the coupled equations above to compare as (e.g., in the free IF2 equation):

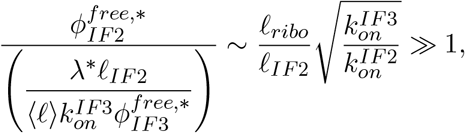

coming from the large size of the ribosome compared to the initiation factors. In addition, the derivative of the coupling terms, which appear in the optimality condition and therefore in identifying the optimal abundances, are all of the form 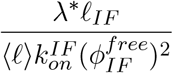 compared to the main term. This scales scales as 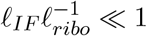 at the self-consistent solution. Hence, neglecting the coupling is justified as an approximate solutions near the optimum, and we obtain for the free concentrations of IFs:

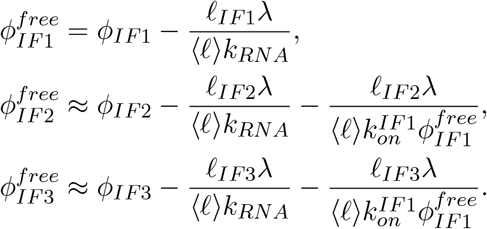

Substituting these in the expression for the initiation time, equation S23, and using the optimality condition (equation 5, we find that no simple solution exist for the non symmetric case of 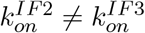. Since the on-rates should be similar for IF2 and IF3 (difference in size should only lead to modest difference in on-rates coefficient, by roughly (*𝓁*_*IF*2_*/𝓁*_*IF*3_)^1*/*3^ ≈ 1.7 assuming Stokes scaling), the symmetric case is approximately correct. We report the symmetric solution for simplicity. The final optimal solutions for the three factors for the sub-scheme solved here is:

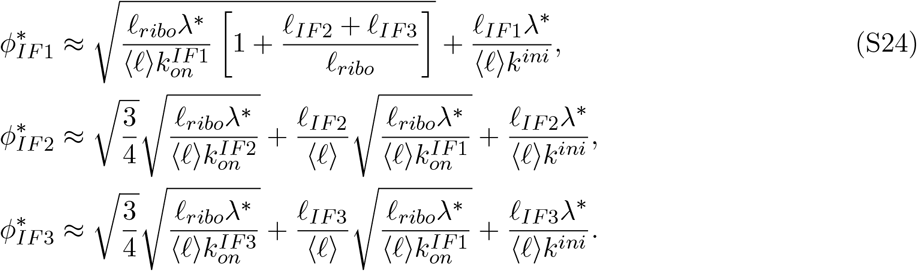

The form of the solution is again similar to that derived for the simpler translation termination scheme (c.f., equation S3), with three differences, each of which has an intuitive interpretation. First, the factor 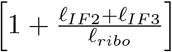 in the IF1 solution arises as a result of IF1 binding being last in our initiation pathway. Indeed, IF1 concentration also influences free IF2 and IF3 concentration, leading to additional selective pressure to increase its abundance. In effect, the molecular species waiting for IF1 to diffuse to its target is not only the ribosome, but the ribosome with IF2 and IF3 bound, and a total amino acid weight *𝓁*_*ribo*_ → *𝓁*_*ribo*_ + *𝓁*_*IF*2_ + *𝓁*_*IF*3_. Second, the factor of 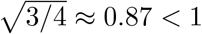 for IF2 and IF3 (corresponding to the symmetric case), arising from the parallel pathway for IF2 and IF3 rendering the process more efficient. We therefore see that the correction from having multiple reactions in parallel is modest (0.87 vs. 1). The third difference to the simpler case of translation termination are the second terms for IF2 and IF3, corresponding to the additional delay incurred by binding of IF1. These come from the assumed sequential nature of our initiation scheme (Fig. S4). In such cases, factors binding earlier have to be present at higher abundances to account for their wait times for later binding events. The exact form of this correction term would be different for more complex assembly pathways (but would be captured by average delays from other factor binding).

### S7.2 Pathway including subunits joining

The solutions above (equations S24) are for the reduced scheme (boxed in Fig. S4). The full solutions includes the delay arising from 50S subunit binding. Including subunit joining requires the solution of an additional equation for the steady-state concentration of species with all three initiation factors, mRNA and initiator tRNA waiting for subunit joining (species *R*_123*m*_ in Fig. S4, denoted *ϕ*_123*m*_ in units of proteome fraction). The equation to solve for *ϕ*_123*m*_ can be obtained from the 50S ribosome subunit conservation equation:

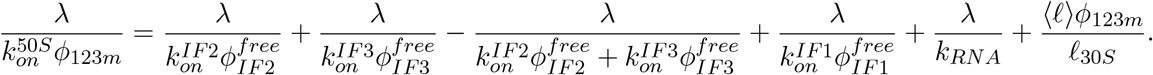

*ϕ*_123*m*_ appears in the equations for the free concentration of the initiation factors (from the conservation equations), and also leads to the appearance of a new term in the expression for the initiation time τ_*ini*_ (equation S23) corresponding to this step: 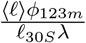.

These two additions, resulting from the parallel branch of 50S joining, can be simplified due to a separation of scales between the various terms. For large initiation factor concentrations, the corresponding mass action terms in the equation for *ϕ*_123*m*_ negligibly contribute to the solution. In this regime, the new term involving *ϕ*_123*m*_ in the initiation time τ_*ini*_ does not alter the form the optimal abundances of IF1, IF2, and IF3 beyond adding a constant term. Hence, in the regime of high free IF concentration, the optimality condition has the same form as derived in the previous section.We can therefore obtain *ϕ*_123*m*_ assuming large IF concentration, denoted 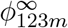:

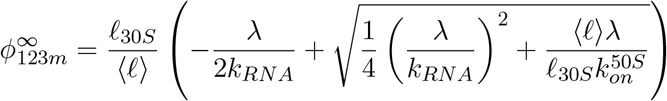

This solution will be self-consistent provided (for all initiation factors):

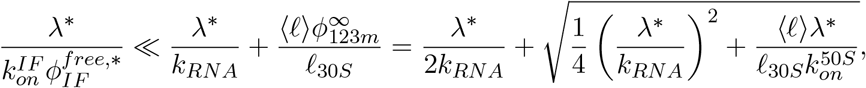

It therefore suffices to show:

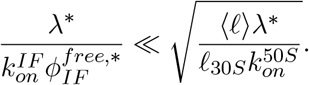

Using our optimality condition on 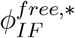 (equation S24) assuming no contribution from *ϕ*_123*m*_ (self-consistency), and converting association rates in units µM^−1^s^−1^, the above condition reduces to:

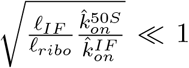

The self-consistency condition is met both because initiation factors are smaller than ribosomes *𝓁*_*IF*_ ≪ *𝓁*_*ribo*_, and because the on-rate for subunit joining is lower than initiation factor binding 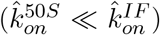, given again the size differences. The solution, including the contribution from ribosome subunits joining is then:

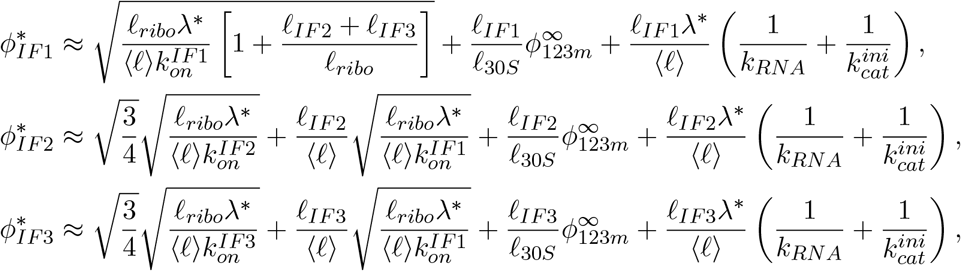

where for *K*_*RNA*_ much faster than the association between the subunits, 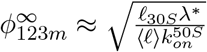.

## S8 Summary of optimal solutions

Solutions for the factor predicted optimal abundances as a function of effective biochemical parameters and the growth rate at the optimum, are presented in Table S3. The table breaks down terms in each solution by categories: direct diffusion term (arising from diffusive search time), catalytic sequestration, and delay incurred by the diffusion of other proteins in part of the cycle of the factor of interest. Solutions are listed in terms of on-rate 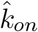 (units of µM^−1^s^−1^). The aaRS solution follows a different form:

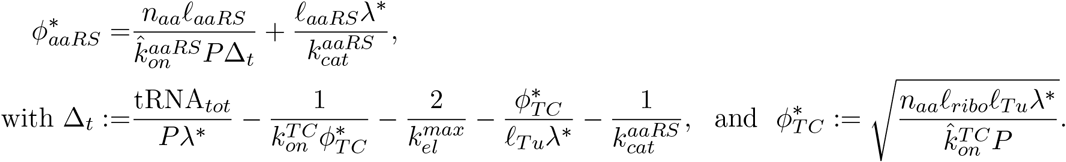

**Table S3:**
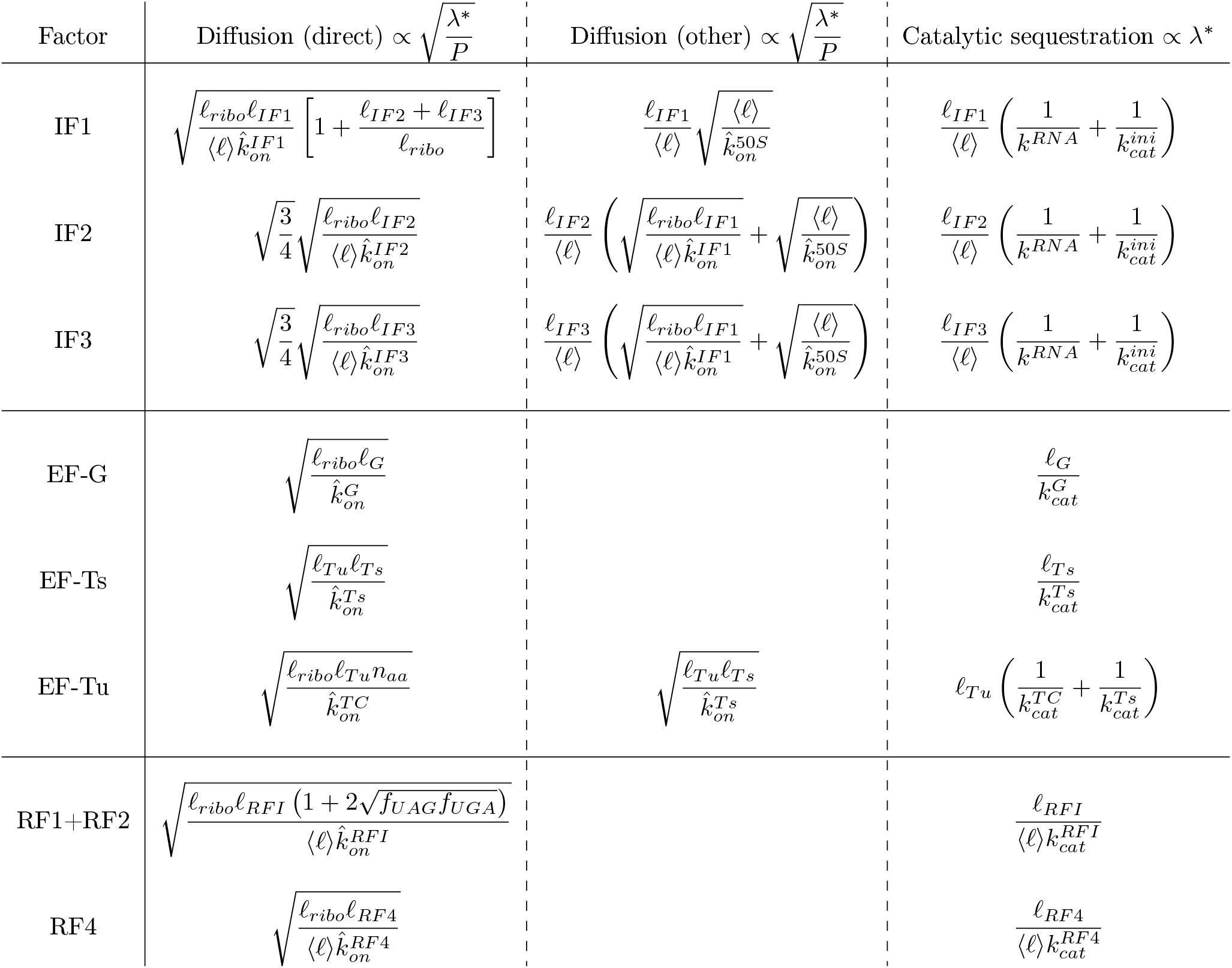
Compilation of predicted optimal abundances for translation factors. The optimal abundance is the sum of the terms in each row. Columns correspond to contributions of different nature (diffusion of factor itself, diffusion of other factors involved in the factor’s cycle, catalytic term). Terms must be multiplied by the common factors indicated in each column’s header (∝).

## S9 Estimation of optimal abundances

To compare prediction from our parsimonious framework (Table S3) requires specific values of kinetic parameters. We use empirical measurements together with scaling relations to estimate these kinetic parameters.

Catalytic rates for many enzymes have been measured *in vitro*, but the obtained values can be sharply incompatible with kinetic parameters that have been measured in the cell. An example is tRNA synthetases. Tallying the measured *k*_*cat*_ for all wild-type *E. coli* aaRSs [24], we find a median value of 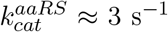, and 80% of reported value below 6 s^−1^. The total molar concentration of aaRSs in the cell is comparable to the total number of ribosomes, and the elongation speed of ribosome is above 15 a.a. s^−1^ [13, 25]. Hence, the absolute minimum catalytic rate to sustain the translation elongation flux needs to obey 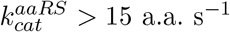, which is much higher than most *in vitro* measured values. To avoid the difficulties in estimating catalytic parameters, and to derive a lower bound on factor abundance from our model, we focus on the diffusive contributions (related to the associate rate) in our predictions, assuming large catalytic rates (*k*_*cat*_ → ∞).

One way to estimate association rates 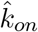 is to assume that the processes are diffusion limited, i.e., following the Smoluchowski relation 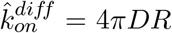, where *D* is the relative diffusion coefficients of the two reactants and *R* the capture radius. Diffusion is a well-characterized process biophysically, and diffusion coefficients of various proteins of different sizes have been measured in the cell [18, 31, 49], including for components of the translation machinery such as ribosomes [4, 59], and tRNAs [55, 67]. Volkov et al report an EF-Tu diffusion coefficient (and a similar measurement for a major diffusive state of tRNAs) of ≈ 3 µm^2^s^−1^. Since ribosomes are nearly immobile, this can be used in an estimate of 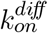, and taking *R* ≈ 10 nm as the rough size of the ternary complex (EF-Tu bound to charged tRNAs), we get in that case for a reaction purely diffusion limited 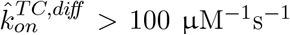. *In vivo* estimates based on kinetic measurements of elongation [13] find 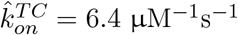, suggesting that the idealized diffusion limited Smoluchowski regime is not applicable in this case. Multiple reasons could explain this, such as rotational diffusion, large off-rate requiring multiple binding events before productive encounter, or possibly because the ternary complex needs to sample multiple non-cognate sites to find a cognate target (thereby slowing its diffusive search).

To circumvent these difficulties, we anchored our association rates to the estimated *in vivo* association rate for the ternary complex, 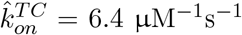 [13], and rescale the association rate by diffusion of related components, i.e., 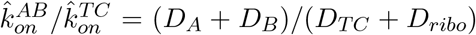, where *D*_*i*_ is the diffusion coefficients for the molecular species *i*. While the *in vivo* diffusion coefficient for a number of component of the translation apparatus exist [4, 55, 59, 67], several factors do not have measured diffusion coefficients. For these, we use the cubic root scaling from the Stokes-Einstein relation [49], see Table S4.

Additional measured quantities required to compute our estimates are: the growth rate λ^*^ = 5.5 × 10^−4^ s^−1^ (21 min doubling time), tRNA concentration (estimated from the tRNA to ribosome ratio of 6.5 [15]), the maximum elongation rate 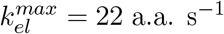 [13], the in-protein amino acid concentration *P* = 2.5 M [11, 30].

Finally, to estimate the ribosome proteome fraction, we subtract from the total concentration to the core translation factor estimated from ribosome profiling (*ϕ*_*R*_ = 0.335) the optimal abundance estimates for the other factors, i.e., 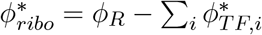. Results displayed in Fig. 4 are listed in Table S5.

**Table S4:** Protein sizes and diffusion coefficients. Unless otherwise noted, protein sizes are taken for *E. coli* [29] ^#^For the ternary complex, the total mass of tRNA+EF-Tu was converted to an equivalent amino acid length. ^†^For aaRS, the proteome fraction weighted (estimated from ribosome profiling [38]) average size (accounting for varying complex stoichiometries) in *E. coli* was taken.

| Factor | Protein size | Diffusion coefficient ($\mu\text{m}^2 \text{s}^{-1}$) |
| --- | --- | --- |
| Ribosome | $\ell_{ribo} = 7336$ [69] | $D_{ribo} = 0.05$[4, 59] |
| 30S subunit | $\ell_{30S} = 3108$ [69] | $D_{subunits} = 0.15$[4, 59] |
| TC | $\ell_{TC} = 630^\#$ | $D_{TC} = 3$ [67] |
| tRNA | | $D_{tRNA} = 8$ [55, 67] |
| IF1 | $\ell_{IF1} = 72$ | $D_{IF1} = D_{TC} \sqrt[3]{\frac{\ell_{TC}}{\ell_{IF1}}}$ |
| IF2 | $\ell_{IF2} = 890$ | $D_{IF2} = D_{TC} \sqrt[3]{\frac{\ell_{TC}}{\ell_{IF2}}}$ |
| IF3 | $\ell_{IF3} = 180$ | $D_{IF3} = D_{TC} \sqrt[3]{\frac{\ell_{TC}}{\ell_{IF3}}}$ |
| EF-G | $\ell_G = 704$ | $D_G = D_{TC} \sqrt[3]{\frac{\ell_{TC}}{\ell_G}}$ |
| EF-Ts | $\ell_{Ts} = 283$ | $D_{Ts} = D_{TC} \sqrt[3]{\frac{\ell_{TC}}{\ell_{Ts}}}$ |
| EF-Tu | $\ell_{Tu} = 394$ | $D_{Tu} = D_{TC} \sqrt[3]{\frac{\ell_{TC}}{\ell_{Tu}}}$ |
| aaRS | $\ell_{aaRS} = 1196^\dagger$ | $D_{aaRS} = D_{TC} \sqrt[3]{\frac{\ell_{TC}}{\ell_{aaRS}}}$ |
| RF1/RF2 | $\ell_{RFI} = 362$ | $D_{RFI} = D_{TC} \sqrt[3]{\frac{\ell_{TC}}{\ell_{RFI}}}$ |
| RF4 | $\ell_{RF4} = 185$ | $D_{RF4} = D_{TC} \sqrt[3]{\frac{\ell_{TC}}{\ell_{RF4}}}$ |

**Table S5:**
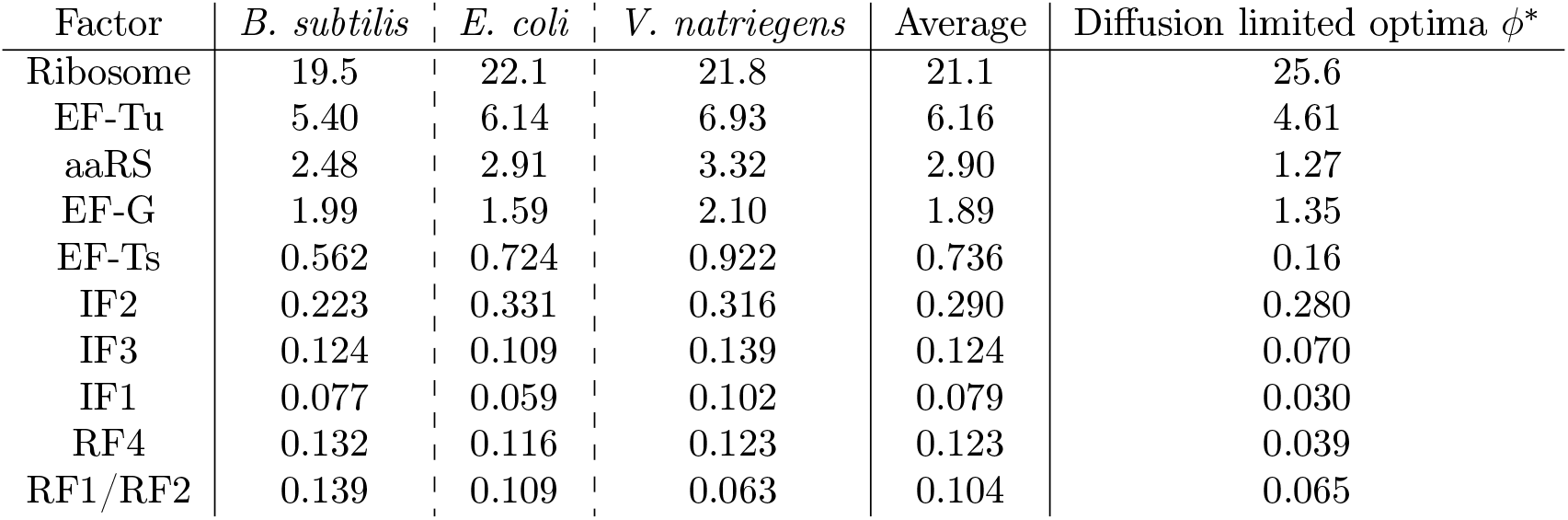
Proteome fraction (in %) of core mRNA translation factors in fast growing *B. subtilis, E. coli*, and *V. natriegens* estimated from ribosome profiling [34, 38] (shown in Fig. 1B), average across the three species, and estimated diffusion-limited optima (Fig. 4).

